# Dominant *Vibrio cholerae* phage exhibits lysis inhibition sensitive to disruption by a defensive phage satellite

**DOI:** 10.1101/790493

**Authors:** Stephanie G. Hays, Kimberley D. Seed

## Abstract

Bacteriophages and their bacterial hosts are locked in a dynamic evolutionary arms race. Phage satellites, selfish genomic islands which exploit both host bacterium and target phage, further complicate the evolutionary fray. One such tripartite system involves the etiological agent of the diarrheal disease cholera – *Vibrio cholerae*, the predominant phage isolated from cholera patients – ICP1, and a phage satellite – PLE. When ICP1 infects *V. cholerae* harboring the integrated PLE genome, PLE accelerates host lysis, spreading the PLE while completely blocking phage production protecting *V. cholerae* at the population level. Here we identify a single PLE gene, *lidI*, sufficient to mediate accelerated lysis during ICP1 infection and demonstrate that LidI functions through disrupting lysis inhibition – an understudied outcome of phage infection when phages vastly outnumber their hosts. This work identifies ICP1-encoded holin and antiholin genes *teaA* and *arrA* respectively, that mediate this first example of lysis inhibition outside the T-even coliphages. Through <u>l</u>ysis <u>i</u>nhibition <u>d</u>isruption, LidI is sufficient to limit the number of progeny phage produced from an infection. Consequently, this disruption bottlenecks ICP1 evolution as probed by recombination and CRISPR-Cas targeting assays. These studies link novel characterization of the classic phenomenon of lysis inhibition with a conserved protein in a dominant phage satellite, highlighting the importance of lysis timing during infection and parasitization, as well as providing insight into the populations, relationships, and evolution of bacteria, phages, and phage satellites in nature.

**Importance:** With increasing awareness of microbiota impacting human health comes intensified examination of, not only bacteria and the bacteriophages that prey upon them, but also the mobile genetic elements (MGEs) that mediate interactions between them. Research is unveiling evolutionary strategies dependent on sensing the milieu: quorum sensing impacts phage infection, phage teamwork overcomes bacterial defenses, and abortive infections sacrifice single cells protecting populations. Yet, the first discovered environmental sensing by phages, known as lysis inhibition (LIN), has only been studied in the limited context of T-even coliphages. Here we characterize LIN in the etiological agent of the diarrheal disease cholera, *Vibrio cholerae*, infected by a phage ubiquitous in clinical samples. Further, we show that a specific MGE, the phage satellite PLE, collapses LIN with a conserved protein during its anti-phage program. The insights gleaned from this work add to our expanding understanding of microbial fitness in natural contexts beyond the canonical bacterial genome and into the realm of antagonistic evolution driven by phages and satellites.

## Introduction

In the 50 years following the discovery of bacteriophages (viruses that infect bacteria informally termed ‘phages’) (1, 2) early geneticists used them to reveal basic biological paradigms including that mutations occur randomly providing the diversity upon which selection acts (3) and DNA is the hereditary substance of life (4). Lytic phages, which lyse the host after hijacking bacterial machinery to produce virions (in contrast to lysogenic phages which can integrate into the host), were of particular interest as model systems; of these, *Escherichia coli*’s T1 through T7 lytic phages rose to prominence (5). Early observable phenotypes in rapid lysing T-even mutants, called r mutants, were leveraged to determine how mutations manifest at the molecular level, how mutagens work, and that nucleic acids are decoded in triplets (6, 7). The r mutant phages produce large plaques with clear edges while plaques of wild type (WT) T-even phages are small with fuzzy edges (8). It was determined that edge fuzziness is due to inhibited lysis of bacterial cells adsorbing additional phage after initial infection. This ‘superinfection’ also stabilizes infected cells as measured by optical density (OD) (9). The phenomenon was termed lysis inhibition (LIN) and failure to lyse during LIN was proposed as significant for two reasons. First, it allows for the production of additional progeny phage resulting in a larger burst. Additionally, it protects progeny phage from adsorbing to cells that are already infected, which are not productive hosts for the secondarily adsorbed phages (Fig. 1A & B) (9). Consequently, LIN is considered an important evolutionary adaptation for phages in environments where host bacteria are scarce but free virions are plentiful (10). While such visually discernable phenotypes enabled early genetic inquiries, the molecular underpinnings of LIN remained unknown until recently and have only been identified and characterized in the T-even coliphages where LIN is mediated by holins and antiholins (11–13). Holins are the first step in canonical holin-endolysin-spanin lysis systems for Gram negative bacteria (14, 15). To briefly summarize the process, holins accumulate in the inner membrane acting as lysis timers, until they reach a critical concentration that triggers depolarization of the membrane, making holes that enable endolysins to access the peptidoglycan. After the endolysins digest the cell wall, spanins mediate the fusion between the inner and outer membranes to complete lysis of the cell. During LIN, antiholins inhibit holin oligomerization and triggering thereby stopping progression towards cell lysis (11–13).

**Fig. 1.**
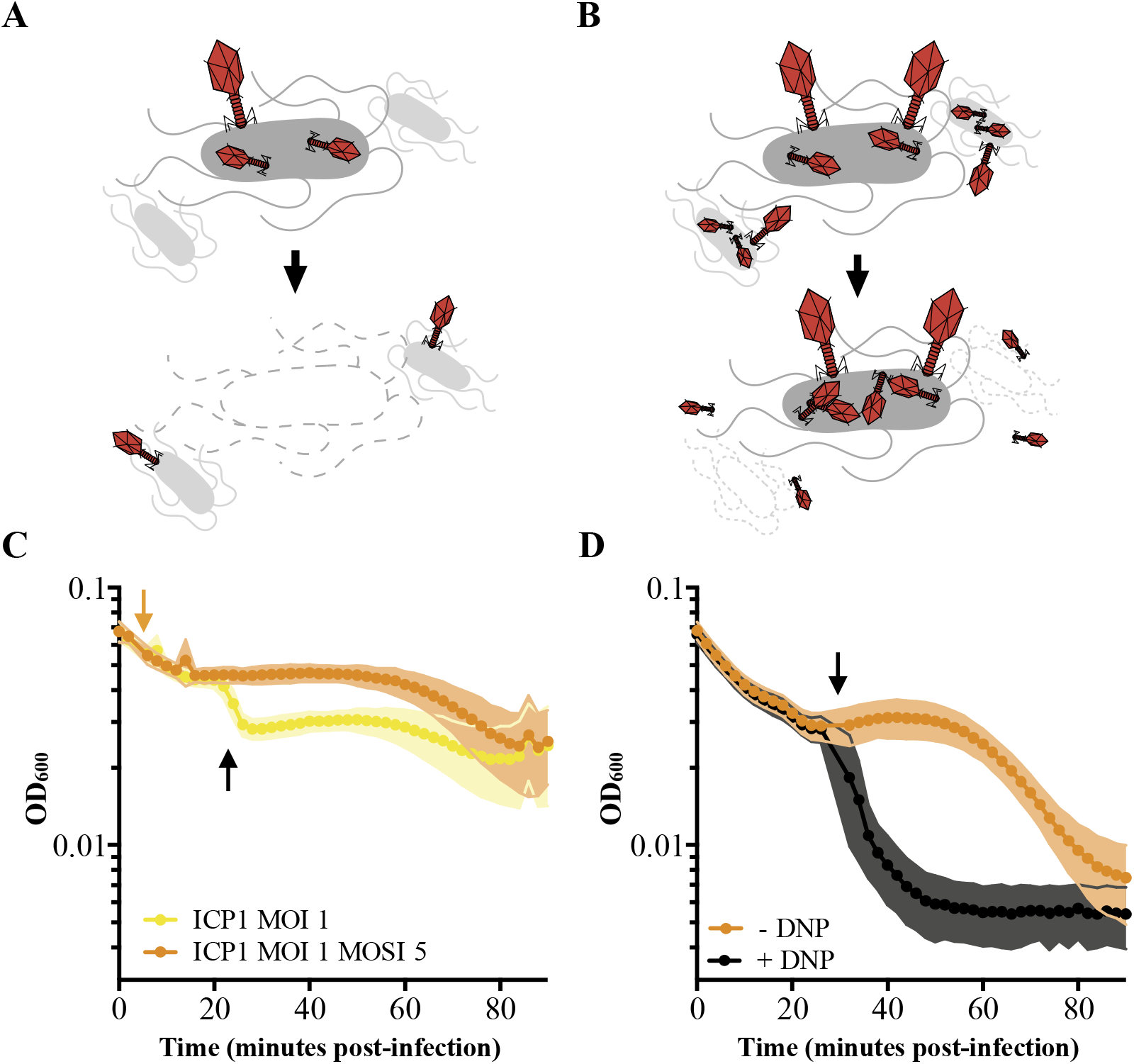
Characterizing LIN. Schematic of T4 infection of *E. coli*. **A** At low MOI, available *E. coli* are readily infected by T4 (red). Under these conditions, an infected cell is hijacked to produce progeny phage and lyses releasing the phages to the environment where they go on to infect neighboring cells. **B** At high MOI, phage outnumber the hosts resulting in initial infection by phage followed by secondary superinfection (second phage infecting the same cell). Superinfection delays lysis through LIN enabling the production of more virions and protecting progeny phage from an environment devoid of uninfected hosts. **C** ICP1 exhibits LIN at intermediate MOIs and upon superinfection. PLE (-) *V. cholerae* infected with ICP1 MOI=1 demonstrate a lysis event 20 minutes post-infection (black arrow) before the OD_600_ stabilizes. Superinfection (orange arrow) of a culture four minutes post infection with ICP1 MOI=1 with ICP1 at a multiplicity of superinfection (MOSI) of 5 triggers LIN and stabilizes the OD_600_ before a lysis event can occur. **D** ICP1 LIN is sensitive to chemical collapse. PLE (-) *V. cholerae* infected with ICP1 MOI=5 maintain OD_600_ for an extended period but the OD_600_ collapses when DNP is added (black arrow).

Lytic phages are abundant in natural environments including marine ecosystems (16) and gut microbiomes (17), making LIN a relevant tool in the molecular arsenal of lytic phages infecting bacteria like *Vibrio cholerae*. *Vibrio cholerae* causes a substantial global health problem as the causative agent of the diarrheal disease cholera (18). Interestingly, in aquatic reservoirs as well as in stool samples from cholera patients, *V. cholerae* co-occurs with phages that target it as a host. The predominant phage in cholera patient samples is ICP1, a lytic myovirus (19). This phage and its host have been locked in a dynamic arms race with no clear winner as both *V. cholerae* and ICP1 continue to be isolated from patients in the cholera endemic region of Bangladesh (19–22). Added into the evolutionary fray is a phage satellite called the PLE (<u>p</u>hage-inducible chromosomal island-like <u>e</u>lement), which is found in the chromosome of *V. cholerae* clinical isolates (23, 24). Previous analysis dating back to 1949 shows that five distinct PLEs with shared genomic architecture appeared over time. Each PLE provides *V. cholerae* with a clear fitness benefit in the fight against ICP1 by completely blocking the production of ICP1 progeny phage (23). PLE hijacks an ICP1-encoded protein to excise from the bacterial chromosome following infection (25), where it replicates to high copy using both PLE and ICP1-encoded proteins (22, 26). PLE is then hypothesized to steal ICP1 structural components to transduce the PLE genome to naïve recipient cells (23). As no infectious ICP1 are produced, PLE activity ultimately defends populations of *V. cholerae* from ICP1 attack, functionally acting as an abortive infection system. In the face of this assault, ICP1 has acquired a functional Type I-F CRISPR-Cas adaptive immune system to target PLE in a sequence-specific manner, restoring ICP1 progeny phage production and overcoming PLE (24, 27). While the full extent of ICP1, PLE, and *V. cholerae* interactions are unknown and continuing to evolve, efforts to understand how the PLE completely restricts ICP1 production have yet to identify any single PLE-encoded gene product that is necessary for inhibition, suggesting that multiple inhibitory strategies act synergistically to fully eliminate phage production (22, 23, 26). From the only characterized processes, namely PLE excision and replication, it is clear that PLE requires phage-encoded products, consequently depending on ICP1 for horizontal transmission, but, uncharacteristic of other phage satellites like the well characterized <u>*S*</u>*taphylococcus* <u>*a*</u>*ureus* <u>p</u>athogenicity islands (SAPIs) which only decrease the number of progeny virions produced, PLE completely abolishes ICP1 production – a balancing act that likely requires the exploitation of select products at just the right time during the 20 minutes before PLE-mediated accelerated lysis.

The accelerated lysis program in *V. cholerae* harboring PLE led us to investigate the prolonged infection in strains without PLE where we discovered that ICP1 exhibits LIN. In this work we report this first characterization of archetypal lysis inhibition outside of *E. coli*, and we characterize ICP1 LIN mechanisms in *V. cholerae* by identifying previously uncharacterized ICP1 genes with holin and antiholin activity, named *teaA* and *arrA* respectively. Subsequently, we also report that PLE-mediated accelerated lysis is phenocopied by collapse of LIN mediated by a single PLE-encoded gene that we call *lidI* for its lysis inhibition <u>d</u>isruption activity. All PLEs encode LidI (a previously undiscovered open reading frame with unknown predicted function), highlighting a conserved strategy to antagonize an aspect of phage lifecycle not previously known to be targeted by parasitic satellites. While LidI normally functions in the context of the PLE, it alone is sufficient to decrease the yield of phages from an infected *V. cholerae* culture imposing an evolutionary bottleneck on phage populations as probed by the potential of ICP1 isolates to recombine for genetic innovation and to escape bacterial CRISPR-Cas.

## Results

### ICP1 causes lysis inhibition (LIN)

After infection at a low multiplicity of infection (MOI), ICP1 completes the production of virions within 20 to 25 minutes after infection of PLE (-) *V. cholerae* (23). This timeframe is markedly abbreviated with respect to the 90 minutes that pass before visible lysis of PLE (-) *V. cholerae* infected with ICP1 at high MOI (23). This incongruity prompted us to test ICP1 infections at intermediate MOI. When infecting PLE (-) *V. cholerae* with ICP1 at MOI=1, we observed an early lysis event 20 minutes post-infection after which the optical density (OD) of the culture stabilized (Fig. 1C). Such lysis kinetics are consistent with canonical LIN by T-even coliphages wherein a portion of infected cells release progeny phages that then trigger LIN – ultimately delaying lysis of the remaining population of cells (9). These similarities led us to hypothesize that ICP1 exhibits LIN in *V. cholerae*.

During canonical LIN in T-even phages, superinfection – the secondary adsorption of phages after initial phage infection – stabilizes the OD of infected *E. coli* cultures as cells stay intact instead of lysing (Fig. 1B). To determine whether superinfection by ICP1 in PLE (-) *V. cholerae* exhibits this same characteristic, we infected cultures with ICP1 at MOI=1, let phage adsorb for four minutes, and then added ICP1 at a multiplicity of superinfection (MOSI) of 5. As expected of a phage exhibiting LIN, the culture OD was stabilized, eliminating the early lysis event normally seen during infection at MOI=1 (Fig. 1C).

Previously characterized LIN in *E. coli* phages is sensitive to changes in membrane proton motive force (PMF) and can be disrupted by the addition of energy poisons. One such poison, the ionophore 2,4-dinitrophenol (DNP), collapses the PMF and subsequently disrupts T2 and T4 LIN, resulting in rapid lysis of infected *E. coli* (28, 29). To test if ICP1 LIN is similarly linked to PMF, we exposed PLE (-) *V. cholerae* infected at high MOI to DNP (Fig. 1D). We observed the expected crash in OD following the addition of DNP, further supporting the conclusion that ICP1 exhibits LIN in *V. cholerae*.

### ICP1 LIN is mediated by the putative holin, *teaA*, and the putative antiholin, *arrA*

Classical lysis by tailed phages employs holins, which accumulate in the inner membrane until they are triggered to form holes prompting cell lysis (Fig. 2A) (30, 31). Though sequenced genomes of ICP1 isolates from cholera patient stool dating back to 2001 are abundant, the molecular basis of ICP1-mediated lysis is not understood. No holin had previously been identified. Inhibition of lesion formation by holins is accomplished by anitholins during LIN in T4 (Fig. 2A) (32), however no antiholins had previously been identified in ICP1 either. To characterize the mechanism underlying ICP1 LIN, we endeavored to find ICP1’s holin and antiholin genes through bioinformatics.

**Fig. 2.**
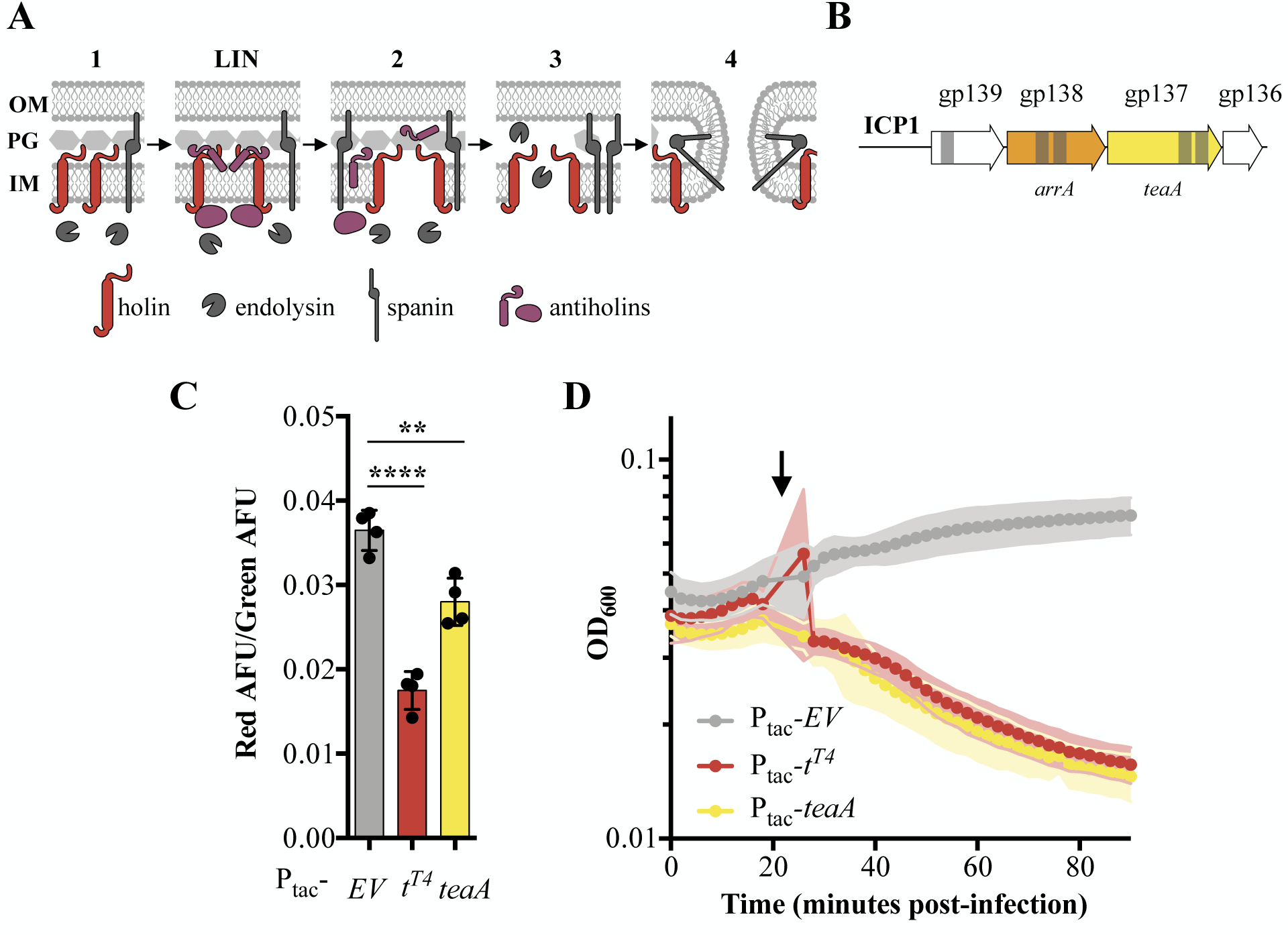
Holin TeaA identification and characterization. **A** Schematic of canonical T4 lysis in *E. coli*. Lysis occurs in four steps with a potential delay caused by LIN. Step one: Phage encoded proteins including holins, endolysins, and spanins accumulate in the cell. In the event of superinfection, antiholins halt the lysis program causing LIN. Step two: Holins are triggered collapsing the PMF and forming holes in the inner membrane (IM) releasing endolysins to the periplasm. Step three: The peptidoglycan is (PG) is degraded by endolysins. Step four: Spanins fuse the inner and outer membrane (OM). **B** TMDs (grey bars) were predicted in three gene products (gp) in a conserved locus of ICP1. **C** Relative PMF state as measured by the ratio of red to green arbitrary fluorescence units from DiOC_2_(3) after ectopic gene expression. Points represent individual replicates; bar height is the average; error bars display the standard deviation of samples. Significance was calculated via one-way ANOVA followed by Dunnett’s test. **** P ≤ 0.0001; ** p ≤ 0.01. **E** Chemical triggering with DNP (arrow) after ectopic gene expression as measured by OD_600_. Points show the average of four replicates; shading shows the standard deviation.

Holins are among the most diverse proteins, however, because they sit in the membrane, all holins include at least one transmembrane domain (TMD) (31). In an effort to find ICP1’s holin, the TMHMM Server v. 2.0 was used to predict TMDs in all of ICP1’s 200+ predicted open reading frames. Further, since holins control lysis timing – an essential component of the phage life cycle – we hypothesized that the holin would be conserved in ICP1 isolates over time. As a result we narrowed our search for the holin down to genes with at least one TMD that were conserved within previously analyzed ICP1 isolates (20). This left three candidate genes in a single cluster within the ICP1 genome (Fig. 2B). Analysis of these ORFs using remote homology and synteny to predict protein function via Phagonaute identified the DUF3154 Pfam (subsequently reclassified as the GTA_holin_3TM (PF11351) family of holins (33–35)) in ICP1 GP137, which we have since named TeaA. Interestingly, not only is TeaA conserved in ICP1 but there are TeaA homologs present throughout marine phage and bacterial genomes suggesting function outside of our phage of interest (Supplemental Fig. 1).

In *E. coli*, canonical holins, including T4’s T-holin, accumulate in the membrane until they reach an adequate concentration to collapse the PMF and form pores or are triggered by an energy poison. Premature triggering of holins results in loss of viability and commits the cell to eventual lysis, decreasing the OD (36, 37). To experimentally test TeaA for holin activity, we exogenously expressed *teaA* in PLE (-) *V. cholerae* in the absence of phage and probed its ability to collapse the PMF and be triggered by the energy poison DNP. We measured PMF using 3,3’-diethloxacarbocyanine iodide (DiOC_2_(3)), a fluorescent green membrane stain that forms red fluorescent aggregates in the presence of intact PMF (38, 39). Upon induction, T-holin^T4^ and TeaA resulted in decreased red fluorescence suggesting holin activity while the empty vector (EV) control did not (Fig. 2C). Holin activity by TeaA was further demonstrated by the rapid decrease in OD after DNP addition, which was comparable to the decrease in OD when DNP was added to T-holin^T4^ expressing *V. cholerae* (Fig. 2D). These similarities to T4’s T-holin motivated the naming of *teaA* for the gene’s <u>T</u>-holin-<u>e</u>sque <u>a</u>ctivity.

Next, we sought to identify an antiholin in ICP1. Antiholins can inhibit pore formation via multiple mechanisms: antiholin TMDs can directly interact with holins in the membrane altering quaternary structure (40, 41), antiholin periplasmic domains can sense signals of superinfection while interacting with holins to block pore formation (42), and cytoplasmic antiholins can bind holins inside the cytoplasm (43). Because antiholins interacting directly with holins within the membrane will contain TMDs and previously characterized periplasmic antiholins contain TMDs related to their tethering to the membrane and subsequent release into the periplasm (44), we continued to focus on the ORFs in ICP1 containing at least one TMD. Given that TeaA is conserved in ICP1 isolates, and lysis timing, which is finetuned by antiholins, is critical to phage fitness, we expected that the antiholin would also be conserved. Antiholins are often found near holin genes, in many cases even utilizing an alternative start site within the holin sequence or occupying an overlapping ORF (45, 46). With these data in mind, we began to investigate GP138, which we subsequently named ArrA, as a potential antiholin because it contains TMDs, is conserved, and is encoded directly next to TeaA (Fig. 2B).

The T4 antiholins, RI^T4^ and RIII^T4^ were initially identified when mutations in these genes demonstrated a rapid lysis plaque morphology. Phages lacking functional antiholins form plaques with sharply defined edges because LIN no longer occurs hence no apparent fuzziness on the periphery of the plaque. In an effort to find similar rapid-lysing ICP1 mutants with antiholin modifications, WT ICP1 was plaqued on CRISPR-CAS (+) *V. cholerae* containing a spacer targeting *arrA*. The resulting plaques had a mixture of plaque edge phenotypes including clear-edged plaques (Fig. 3A). To determine that these putative mutant LIN ICP1 were deficient in ArrA, we supplied ArrA *in trans* and recovered the fuzzy edge phenotype (Fig. 3B). Because the CRISPR-Cas targeting yielded a mixed pool of mutants, we engineered a clean Δ*arrA* ICP1 strain in order to further confirm ArrA function. Of note, and consistent with *arrA* acting as an antiholin, we could successfully construct a clean knockout of *arrA* suggesting that, despite its conservation, it is not an essential ICP1 gene. We hypothesized that if ArrA is an antiholin, Δ*arrA* ICP1 would have characteristically clear-edged plaques. Indeed, while the Δ*arrA* mutant successfully forms plaques, it displays the expected clear-edged plaque phenotype suggesting that ArrA is an <u>a</u>ntiholin whose <u>a</u>bsence results in rapid lysis motivating its name (Fig. 3C).

**Fig. 3.**
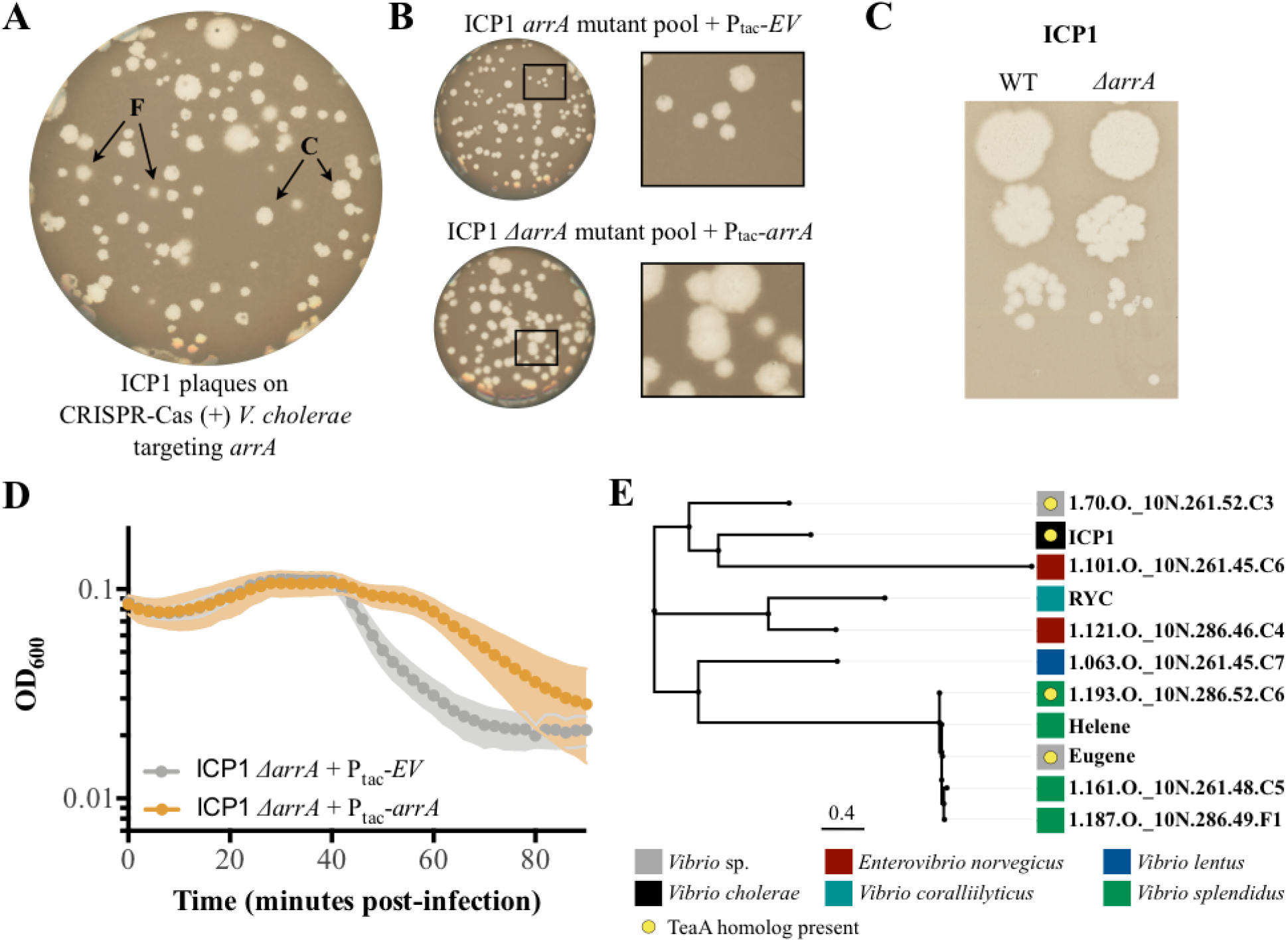
Antiholin ArrA identification and characterization. **A** Initial plaquing of WT ICP1 on CRISPR-CAS (+) *V. cholerae* targeting *arrA* yielded a mixture of plaque phenotypes. F indicates fuzzy-edged plaques. C indicates clear-edged plaques. **B** Plaques with clear edges can be found in populations of phage that overcame targeting of *arrA* with various mutations in *arrA* when plaqued on *V. cholerae* harboring an EV control (Top). Providing ArrA *in trans* restores the fuzzy edge phenotypes within the mutant population. **C** Using a repair template we created a clean *arrA* deletion and found that Δ*arrA* ICP1 yield plaques with clear edges. **D** During infection, once lysis begins, Δ*arrA* ICP1 causes rapid lysis consistent with abolishment of LIN. When ArrA is expressed *in trans* the LIN phenotype is rescued: a small lysis event is visible 40 minutes post-infection after which the OD is stabilized for about 30 minutes. **E** PBLAST was used to find proteins with 20% identity to ArrA over 75% of the query; these homologs are displayed in an unrooted tree displaying the name of the phage the protein is found in. Colored blocks show the host each phage infects, and yellow circles denote that a protein with homology to TeaA is present in the phage.

In addition to changing plaque morphology, the T4 antiholin mutants demonstrate accelerated lysis kinetics in liquid culture (11, 13), motivating us to test infection with ICP1 Δ*arrA* in liquid cultures. Attempts at making high titer stocks of Δ*arrA* ICP1 were unsuccessful (consistent with decreased phage yields due to a lack of LIN) hindering our ability to test infection at high MOI. Instead, we infected PLE (-) *V. cholerae* with Δ*arrA* ICP1, waited for approximately two cycles of infection to complete, and then, once lysis began, we observed a rapid crash in culture OD (Fig. 3D) in contrast to the more prolonged decline seen in WT infections exhibiting LIN (Fig. 1C). To be sure this was due to ArrA, we supplied ArrA *in trans* and recovered lysis kinetics characteristic of LIN: instead of the rapid lysis seen in cultures with the mutant phage without complementation, we see a stabilization of OD for an additional 30 minutes before the culture clears (Fig. 3D) further supporting the conclusion that ArrA is an ICP1-encoded antiholin that helps to regulate lysis timing.

Interestingly, BLASTP analysis of ArrA yielded fewer homologs than TeaA (Supplemental Fig. 1), however, when we used less stringent search parameters we found that some organisms containing TeaA homologs also contain ArrA homologs (Fig. 3E). This suggests that there are potential homologous LIN systems – complete with both holin and antiholin – present in other phages outside of ICP1. In contrast, the presence of ArrA antiholin homologs without TeaA holin homologs raises the question: what are antiholins doing on their own? Perhaps they have evolved functionality with holins divergent enough to no longer be considered homologous to TeaA under our search parameters, they had previous functions before evolving to moderate lysis timing, or they have been coopted for divergent functions much like holins have been (47, 48).

### PLE accelerates ICP1-mediated lysis

Thus far, all characterization of ICP1 LIN was completed in PLE (-) *V. cholerae.* In clinical samples of *V. cholerae*, an increasing percentage of collected isolates from Bangladesh harbor PLE (21). PLE is a phage satellite, and by definition, a parasite of ICP1; although the mechanisms that PLE deploys to inhibit and hijack ICP1 are not completely understood, the lysis kinetics for high MOI ICP1 infections of *V. cholerae* vary depending on the presence of PLE. Consistent with previous experiments (23), ICP1 infection of PLE (-) *V. cholerae* at high MOI results in a gradual lysis which reaches the lowest OD around 90 minutes after the initial addition of phage. In contrast, upon infection of *V. cholerae* containing the integrated PLE 1 genome, rapid lysis is observed starting 20 minutes post infection (Fig. 4A). This acceleration in lysis timing could conceivably be caused by a number of processes: PLE deploying its own lysis machinery, PLE modulating the expression or stability of components of ICP1’s lysis machinery, PLE collapsing LIN, or any combination of these mechanisms could force infected PLE 1 *V. cholerae* to lyse earlier than PLE (-) *V. cholerae*.

**Figure 4.**
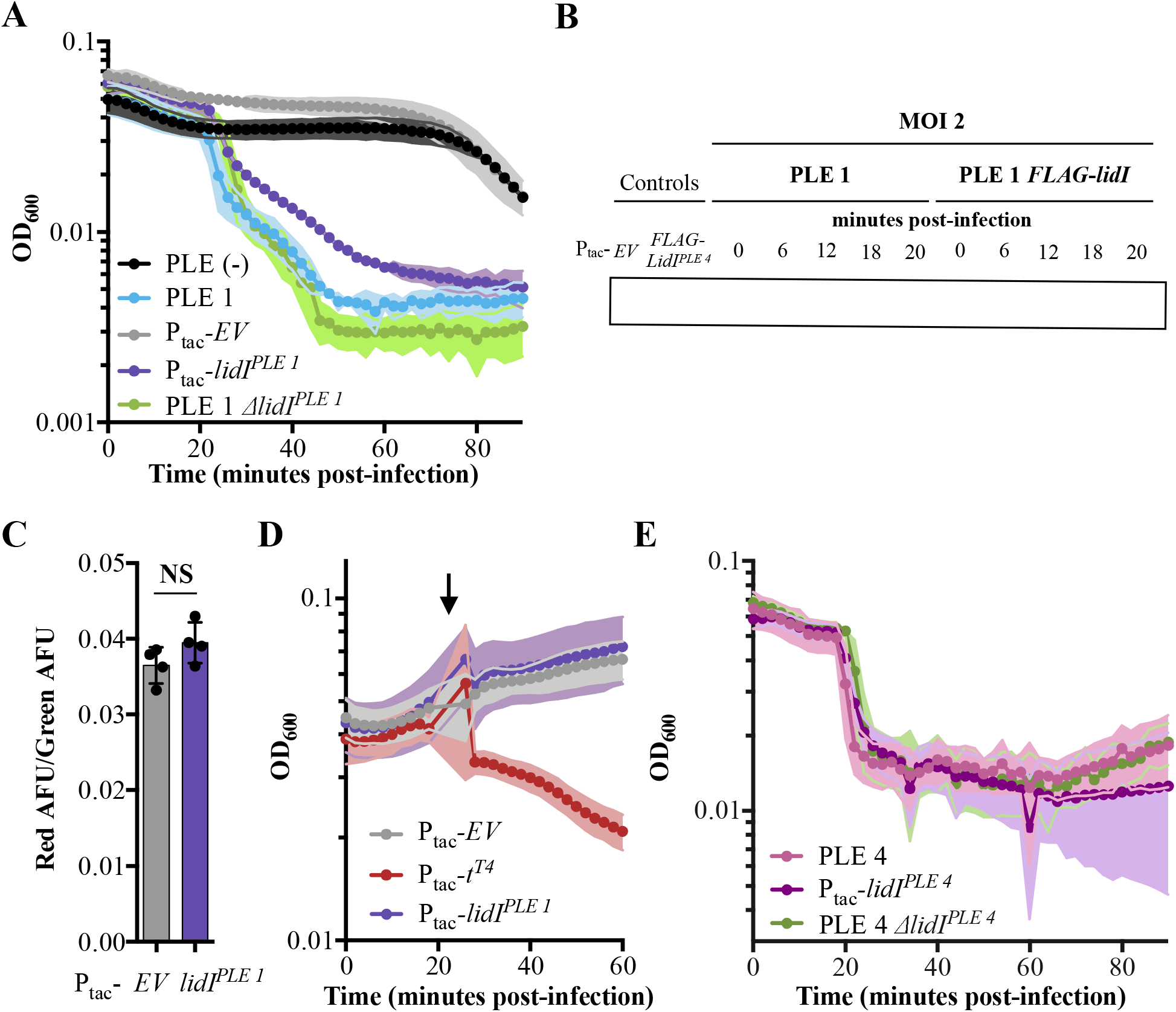
Accelerated lysis by PLE and *lidI*^*PLE 1*^. **A** The OD_600_ of ICP1 MOI=5 infections was followed in PLE (-), PLE 1, and PLE 1 Δ*lidI V. cholerae* strains as well as *V. cholerae* strains containing induced *EV* and *lidI*^*PLE 1*^ constructs. Data points represent the average reading of n≥3 replicates and shaded regions display the standard deviation of experiments. **B** Tagged *lidI*^*PLE1*^ expressed *in trans* is readily observable by Western blot. Tagged *lidI*^*PLE 1*^ in the native PLE context is visible 18 to 20 minutes post infection with ICP1 MOI=2. Complete blot available in Supplemental Fig. 3. **C** PMF state as measured by the ratio of red to green arbitrary fluorescence units from DiOC_2_(3) after ectopic gene expression. NS signifies ‘no significance’. **E** Chemical triggering with DNP (arrow) after ectopic gene expression as measured by OD_600_. Points show the average of four replicates; shading shows the standard deviation. **F** The OD_600_ of ICP1 MOI=5 infections was followed in PLE (-), PLE 4, and PLE 4 Δ*lidI V. cholerae* strains as well as *V. cholerae* strains containing induced *EV* and *lidI*^*PLE 4*^ constructs. Data points represent the average reading of n≥5 replicates and shaded regions display the standard deviation of experiments.

To start investigating the mechanistic underpinnings of accelerated lysis in the presence of PLE 1, we scrutinized the PLE for ORFs encoding potential lysis machinery. The PLE 1 genome contains 25 previously annotated ORFs of which only two (the integrase, *int*, (25) and a replication initiation factor, *repA*, (26)) have characterized functions, prompting us to evaluate if any of the remaining ORFs might include a holin. Initial analysis using the TMHMM Server v. 2.0 (49) revealed no TMDs in PLE 1 ORFs, potentially ruling out the presence of PLE-encoded lysis machinery. Previous studies revealed that all five known PLEs demonstrate accelerated lysis (23) so we next turned to the genomes of the other four PLEs which contain up to 29 open reading frames. The two earliest known PLEs, (PLE 4 and PLE 5) contain two ORFs with predicted transmembrane domains: ORF 2 and ORF 26 in both PLEs. ORF 2 does not have homologs in PLE 1 or PLE 2, suggesting that it is not a conserved player in the accelerated lysis phenotype. Consequently, we focused on ORF 26. Though no homologs were immediately obvious outside of PLE 4 and PLE 5, the conservation between PLEs suggested the presence of previously unannotated ORFs. Through further analysis, ORF 24.1^PLE 3^, ORF 24.1^PLE 2^, and ORF 20.1^PLE 1^ were discovered, which are homologous to ORF 26 in PLE 4 and PLE 5 and contain transmembrane domains. We subsequently named these genes *lidI* and the homologs cluster neatly into two groups: *lidI*^*PLE 1*^ and *lidI^PLE 2^* are short 66 amino acid predicted proteins with four synonymous SNPs between them, while *lidI^PLE 3^*, *lidI*^*PLE 4*^ and *lidI^PLE 5^* encode larger proteins at 121 amino acids in length with three SNPs that result in two amino acid substitutions (Supplemental Fig. 2). These ORFs have no significant homology to other genes and no predicted functional domains beyond their shared TMDs.

Next, to confirm the expression of the newly discovered *lidI* genes, we tagged LidI^PLE 1^ in the native PLE context and evaluated its expression prior to and during ICP1 infection (Fig 4B & Supplemental Fig. 3). We could not visualize FLAG-LidI^PLE 1^ in the absence of phage infection, however, when we infected with ICP1 at MOI=2, FLAG-LidI^PLE 1^ was detectable by Western blot. During the course of infection, which lasts only 20 minutes in PLE (+) *V. cholerae*, FLAG-LidI^PLE 1^ was only detectable late in infection – 18 to 20 minutes post phage addition and immediately prior to the sudden decrease in OD characteristic of PLE-mediated accelerated lysis (Fig. 4B & Supplemental Fig. 3).

After confirming that *lidI*^*PLE 1*^ is expressed during infection, we next sought to characterize LidI^PLE 1^ function by exogenously expressing *lidI*^*PLE 1*^ in PLE (-) *V. cholerae.* During infection with ICP1, *lidI*^*PLE 1*^ was sufficient to recapitulate the PLE-mediated accelerated lysis phenotype (Fig. 4A). This is consistent with *lidI*^*PLE 1*^ encoding a holin; however, expression of *lidI*^*PLE 1*^ to the same level in the absence of phage did not alter cellular PMF or make cells susceptible to DNP induced lysis (Fig. 4C&D). These data suggest that *lidI*^*PLE 1*^ does not act as a canonical holin when expressed alone and that its lysis function is dependent on the presence of ICP1.

Having demonstrated that LidI^PLE 1^ is sufficient to recapitulate PLE-mediated accelerated lysis, we wanted to determine if it was also necessary for this phenotype. Interestingly, however, when we knocked out *lidI*^*PLE 1*^ from PLE 1 *V. cholerae* we found that the lysis kinetics were unchanged (Fig. 4A). This finding seemed to be at odds with the conservation of *lidI* homologs in all the known PLEs, therefore we tested *lidI*^*PLE 4*^, a representative of the other cluster of homologs, for conserved function using the same assays. Indeed, *lidI ^PLE 4^* is sufficient to cause accelerated lysis in the absence of PLE, however again we found that it is not necessary – PLE 4 Δ*lidI V. cholerae* strains still exhibit accelerated lysis during high MOI infections (Fig. 4E). Though this was counter to what we expected, timing is of paramount importance to phage fitness (50), often finetuned down to the minute, so it is perhaps unsurprising that the PLE, as a phage satellite, would redundantly encode a means to execute lysis for its own selfish benefit. In an effort to identify other PLE-encoded factors sufficient to recapitulate PLE-mediated accelerated lysis, we cloned each of the PLE 1 ORFs individually into inducible constructs and challenged them with phage. This screen did not successfully identify any other single genes sufficient to accelerate lysis, however, it is important to note that cryptic genes could be responsible, other accelerated lysis systems could require more than one ORF to function, or the timing and expression level of genes expressed outside the context of PLE could be obscuring gene functionality.

### LidI^PLE 1^ functions through lysis inhibition disruption

Discovering that *lidI*^*PLE 1*^ is sufficient for the accelerated lysis phenotype, but that it doesn’t phenocopy what is expected for a holin, we wanted to determine the mechanism for LidI^PLE 1^-mediated accelerated lysis. Since ICP1 exhibits LIN during high MOI infections, we hypothesized that LidI^PLE 1^ causes accelerated lysis by disrupting LIN. LIN occurs when phages outnumber hosts, so changing the multiplicity of infection changes the onset of LIN – at low MOIs, a small fraction of cells produce a burst of phage which is adsorbed by neighbors and this process can repeat a number of times until the majority of host cells are infected and superinfection triggers LIN (Fig. 1A). This is evidenced by the stabilization of OD until complete lysis around 80 minutes in PLE (-) *V. cholerae* strains infected at various MOIs (Fig. 5A). The differential onset of LIN means that accelerated lysis mediated by *lidI*^*PLE 1*^ would also be expected to change in accordance with MOI if it functions by disrupting LIN. Indeed, we observed differential lysis timing dependent on MOI in cultures expressing *lidI*^*PLE 1*^ with up to a 40 minute delay at the lowest MOI (Fig. 5A), suggesting that *lidI*^*PLE 1*^ functions through lysis inhibition <u>d</u>isruption giving the gene its name.

**Figure 5.**
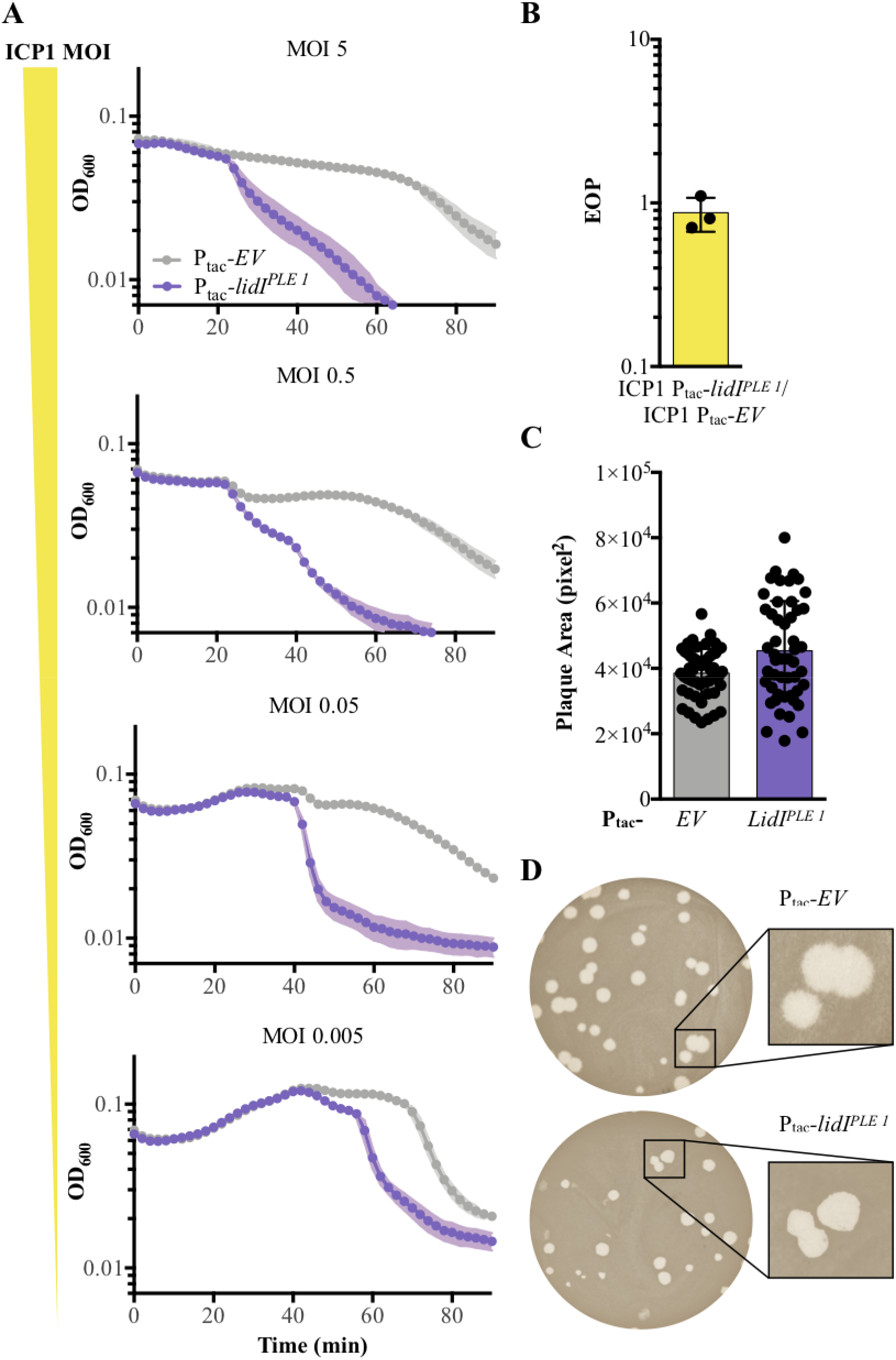
LidI functions through lysis inhibition disruption. **A** The stabilization of OD_600_ due to LIN and subsequent lysis after LIN finishes is dependent on the initial MOI which we observe in infections of EV PLE (-) *V. cholerae*. Consequently, the timing of LidI^PLE 1^-mediated accelerated lysis in *lidI*^*PLE 1*^ PLE (-) *V. cholerae* is also dependent on initial starting MOI. Data points represent the average reading of n=3 biological replicates and shaded regions display the standard deviation of experiments. **B** EOP of induced LidI^PLE 1^ constructs in comparison to EV. Points represent individual replicates; bar height is the average; error bars display the standard deviation of samples. **C** Plaque area was determined by measuring the average number of pixels per plaque of clearly differentiated plaques from three biological replicates of induced *lidI*^*PLE 1*^ and EV containing PLE (-) *V. cholerae.* Points represent the area of individual plaques; bar height is the average plaque area; error bars display the standard deviation of samples. **D** Plaque edge phenotypes were determined for ICP1 plaques on EV and *lidI*^*PLE 1*^ PLE (-) *V. cholerae*.

In line with *lidI*^*PLE 1*^ functioning through collapsing LIN, *lidI*^*PLE 1*^ expression in PLE (-) *V. cholerae* does not change the efficiency of plaquing (EOP) by ICP1 (Fig. 5B). EOP probes the number of successful initial infections at a low MOI so it would not be expected to be altered by the presence or absence of LIN. However, LIN could change the size of plaques as plaque formation involves high MOI infections where the phage are clearing the bacterial lawn to form the plaque. We observed that *lidI*^*PLE 1*^ expression alters this distribution of plaque sizes as well as changes the plaque edge morphology (Fig. 5C&D). Canonical rapid lysis mutations in coliphages show increased plaque size relative to phages capable of LIN which is, similarly, what we would expect from a phage defense mechanism that collapses LIN. However, this is not immediately obvious during expression of *lidI*^*PLE 1*^; instead the distribution of plaque size changes – smalls plaques still exist however there are fewer of them while the average size of the plaques increases (Fig. 5C). While plaque size is one way to measure phage fitness, it is hard to determine exactly what kind of fitness is being measured as multiple variables may impact the end result: plaques may shrink due to decreased burst size from a lack of LIN, increase due to faster lysis kinetics, and be otherwise altered by the presence of clear opposed to fuzzy edges which impact electronic quantification of plaque size. Because of these caveats, the clearest phenotypic change in plaque morphology to indicate disruption of LIN is the loss of fuzzy plaque edges, which we see in PLE (-) *V. cholerae* expressing LidI^PLE 1^ *in trans* (Fig. 5D).

### LidI lowers phage yield

There are varied anti-phage mechanisms in bacteria including defense at the level of phage attachment via receptor modulation, superinfection exclusion, genome degradation by restriction modification and CRISPR-Cas systems, altering fundamental replication, transcription, and translation processes in the phage life cycle, hijacking phage assembly and structural proteins for spread of phage defense and virulence genes, and abortive infections that kill an infected cell protecting the population of uninfected neighbors (51). PLE functionally results in an abortive infection: ICP1 infection of PLE 1 *V. cholerae* results in the lysis of infected cells completely blocking the production of viable phage. Because LIN in T-even coliphages functions to increase phage burst size (9), we hypothesized that LidI^PLE 1^ collapsing LIN could inhibit ICP1 by decreasing progeny phage yield from an infection, though, to our knowledge, no other anti-phage mechanism has been shown to act during the LIN stage of the phage life cycle.

To determine if *lidI*^*PLE 1*^ alone can impact the number of phage produced from an infection, expression was induced prior to, at the time of, and at various intervals after infection with ICP1 MOI=5. Excess un-adsorbed phage were washed off, and, after 100 minutes, cultures were mechanically lysed and the phage yield was determined. Induction of *lidI*^*PLE 1*^ at the time of infection or 20 minutes before infection results in decreased phage yield by one or two orders of magnitude, respectively, in comparison to EV strains (Fig. 6A). Not only did we find this to be true of LidI^PLE 1^, but we also tested the LidI^PLE 4^ homolog for conserved inhibition of ICP1 and observed the same decreased phage yield (Fig. 6B). These data reveal LidI as the first PLE ORF to singlehandedly negatively impact ICP1 phage yield, and it does so by attacking a phage process that no resistance mechanism or phage satellite has been reported to attack: LIN.

**Fig. 6.**
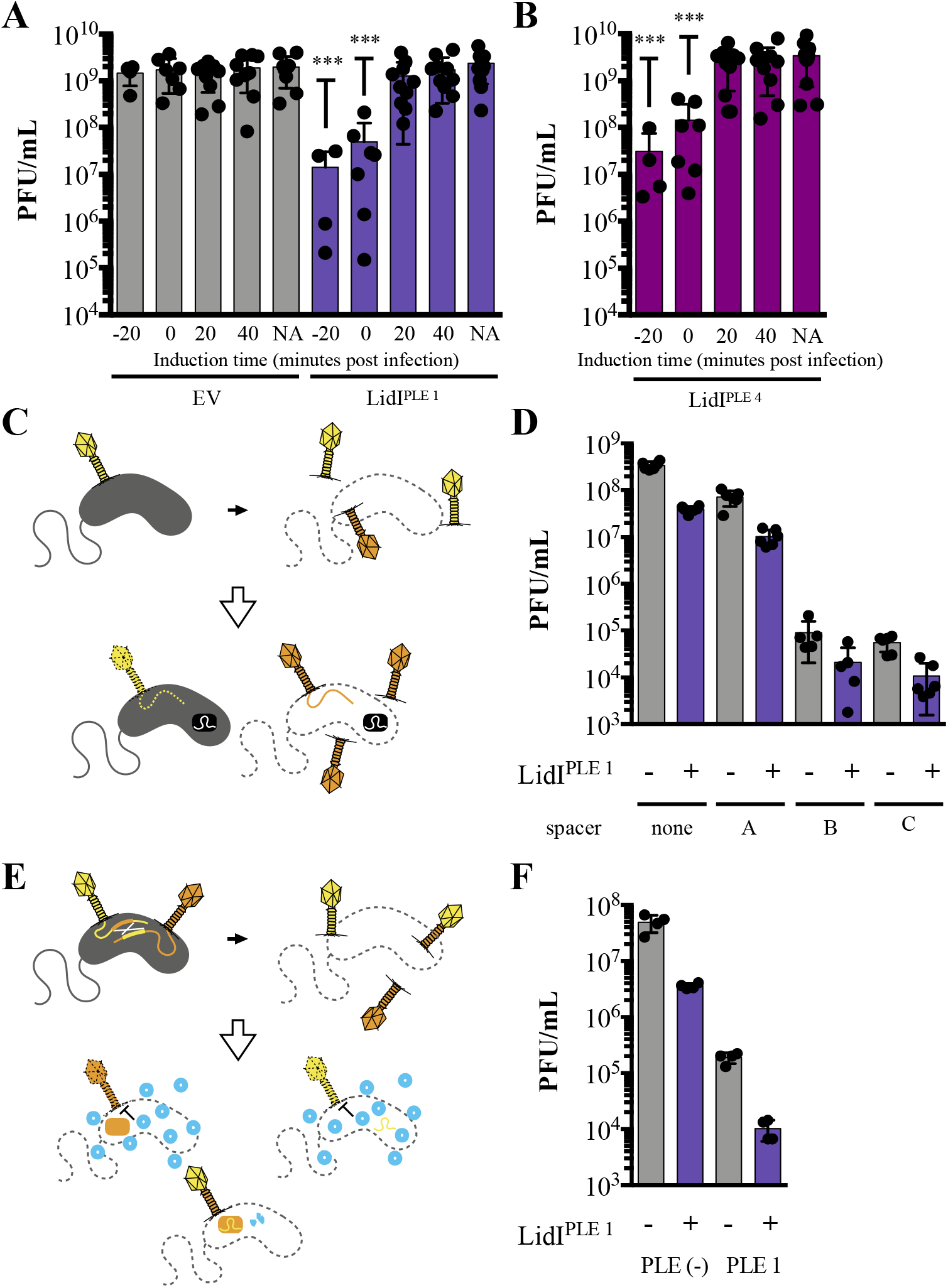
LidI puts a bottleneck on phage populations. **A & B** Phage infection yields were determined from *V. cholerae* with *EV*, *lidI*^*PLE 1*^, and *lidI*^*PLE 4*^ constructs induced at the specified time in respect to phage infection or left uninduced (NA). All samples were mechanically lysed 100 minutes after infection and titers were compared via one-way ANOVA followed by Dunnett’s multiple comparison test comparing all values to titers from the *EV* cultures. Only significant differences are noted. **C** Schematic of evolution as probed by overcoming CRISPR targeting. Top: cultures with induced *EV* and *lidI*^*PLE 1*^ constructs are infected with ICP1 and allowed to produce progeny phage before mechanical lysis. The majority of phages will be WT (yellow) while some phages will harbor mutations (orange). Bottom: The phage populations were then plaqued on *V. cholerae* strains with a CRISPR-Cas system (black rectangle) and different spacers. WT phage infections do not result in plaques because the injected phage DNA is degraded (dotted line) while mutants can overcome the targeting, produce progeny, and lyse the cells. **D** Quantification of the number of phages that could overcome targeting by CRISPR-Cas from populations of progeny phages with and without LidI^PLE 1^. **E** Schematic of evolution as by successful homologous recombination between coinfecting phages. Top: cultures with induced *EV* and *lidI*^*PLE 1*^ constructs are coinfected with ICP1 harboring mutated, non-functional CRISPR-Cas systems: CRISPR*-Cas ICP1 (orange) and CRISPR-Cas* ICP1 (yellow). Bottom: The progeny phage from these infections were the plated on PLE 1 *V. cholerae.* PLE (blue circles) inhibit phage production by all phages that did not successfully recombine to restore a functional CRISPR-Cas system (two-colored phage). **F** Progeny phage from hosts with and without LidI^PLE 1^ co-infected with CRISPR*-Cas ICP1 and CRISPR-Cas* ICP1 were harvested via mechanical lysis. These were then plaqued on permissive (PLE (-) *V. cholerae*) and restrictive (PLE 1 *V. cholerae*) hosts. Significance was determined via one-tailed t-tests between EV and LidI-expressing hosts. For all graphs, data points represent individual values, bar height represents the average value, and error bars represent the standard deviation. * P ≤ 0.05, ** P ≤ 0.01, *** P ≤ 0.001, **** P ≤ 0.0001

From an evolutionary perspective, producing fewer virions equates to less diversity and would be expected to limit the phage’s ability to evolve counterattacks to anti-phage mechanisms or escape through mutation. We hypothesize that PLE-mediated accelerated lysis decreases the ability of ICP1 to evolve in the face of PLE. However, because accelerated lysis is redundantly encoded it is not currently possible to test the impact of delayed lysis on ICP1 evolution in the context of the PLE. We can, however, interrogate how decreasing phage production in the presence of *lidI* in PLE (-) *V. cholerae* puts a bottleneck on ICP1 evolution. To test if LidI^PLE 1^-mediated accelerated lysis is enough to impact diversity through the acquisition of random mutations in the resulting progeny phage population, we exposed PLE (-) *V. cholerae* with and without *lidI*^*PLE 1*^ to ICP1, collected the population of progeny phage, and looked for plaque formation on PLE (-) *V. cholerae* encoding a Type I-E CRISPR-Cas system endogenous to *V. cholerae* classical biotype strains engineered to harbor various anti-ICP1 spacers (Fig. 6C) (27). We see that for each spacer, fewer phages are able to overcome targeting in the phage progeny from *lidI^PLE 1^ V. cholerae* infections than the infections of strains without *lidI*^*PLE 1*^ (Fig. 6D). This effect is due to the LidI-mediated decrease in the population of phages as the frequency of phage escaping stays the same (e.g. ~2 out of every thousand phages can overcome spacer C (Fig. 6D), however because the *lidI^PLE^* ^1^ *V. cholerae* produced an order of magnitude less phage than EV *V. cholerae*, an order of magnitude fewer phages can overcome the spacer in the population; Supplemental Fig 4). While exposure to the CRISPR-Cas system was meant to probe evolution at the level of individual random mutations and proclivity to overcome targeting, phages also readily use homologous recombination to evolve during co-infection. To test the impact of LidI^PLE 1^ on this aspect of evolvability, two variants of ICP1, one devoid of spacers against ICP1 with an inactive Cas1 preventing spacer acquisition (CRISPR*-Cas ICP1) (21) and the other lacking Cas2-3 (CRISPR-Cas* ICP1), were used to coinfect *V. cholerae* strains with and without LidI^PLE 1^. After lysis, progeny phage were tested for their ability to plaque on PLE 1 *V. cholerae*, which is only possible when homologous recombination between the two variants of ICP1 occurred restoring a functional CRISPR-Cas system able to target PLE 1 (Fig. 6E). Predictably, the presence of LidI^PLE 1^ decreased the number of progeny phages per infection that successfully recombined to reconstitute the CRISPR-Cas system and overcome PLE (Fig. 6F; Supplemental Fig. 4). These data suggest that LidI^PLE 1^ puts a bottleneck on the phage progeny population size by accelerating lysis, resulting in decreased diversity while not directly impacting ICP1’s evolvability within an infected cell.

## Discussion

There is increasing awareness of the impact microbiota have on all aspects of life; the carbon fixation by phototrophs in the oceans (52, 53), the health of bees that pollinate plants (54), the production and safety of food (55), and various aspects of human health among others (56). Along with this increased appreciation of microbiomes comes a richer understanding of the roles parasites play; viruses infect hosts throughout the tree of life (57), with bacteriophages, recently re-emerging as a potential tool in biotechnology and the treatment of human disease. Estimates of bacteriophages found in nature are as high as ten to every one bacterium resulting in a dynamic evolutionary arms races between these parasites and hosts where interactions between the adversaries are often mediated by MGEs (58–61). While considering these microbial communities, we must examine not only the organisms present but also the environments in which the arms races occur.

Different environments are battlegrounds that grant different advantages. Considering the tripartite *V. cholerae*/ICP1/PLE system studied in this work, two environments are of particular importance: aquatic reservoirs and the human gut. Aquatic environments seem to be poor arenas for productive infection by vibriophages (62) while harboring dormant, viable but nonculturable *V. cholerae* (63) in contrast to the human gut where *V. cholerae* proliferates to high densities causing disease and being preyed upon by phages (64).

Beyond these vastly different physical environments, there is evidence that the number of organisms present also impacts the arms race between phages and bacteria. Research continues to unveil evolutionary strategies dependent on these ratios. Anti-CRISPRs, the next innovation in counter adaption between phages and bacteria encoding CRISPR-Cas systems targeting those phages, were shown to work in a phage-concentration dependent manner. In short, anti-CRISPRs from a single phage may not necessarily be enough to overcome host defenses, however, if the environment is full of free phages, subsequent phages benefit from the anti-CRISPRs expended in the attempts at infection by previous phages (65, 66). In a similar measurement of microbial populations in an environment, it was recently uncovered that phages can ‘listen in’ on host quorum sensing to influence the decision between lysogeny and lysis depending on the density of host bacteria (67). Both of these discoveries parallel the classically studied LIN system in T-even coliphages in different ways – the anti-CRISPR system overcomes host defenses via multiple subsequent phage infections similar to the multiple phage infections that signal phages should make the most of the host cell before lysing it in the LIN system, and eaves dropping on the host quorum system is akin to LIN because it provides a measure of available hosts. These mechanisms highlight the importance of phages tuning their infection parameters depending on host density and availability. Our discovery of LIN in *V. cholerae* induced by infection with a clinically relevant phage reinforces how important it is for bacteriophages to sense environments with respect to population size. Following ingestion, *V. cholerae* colonizes and blooms in the small intestine before being shed in stool, then going on to further contaminate aquatic reservoirs and promote subsequent ingestion and infection cycles. If large numbers of ICP1 are co-ingested with small numbers of *V. cholerae*, it would be theoretically beneficial for ICP1 to demonstrate LIN in infected cells, to bide time inside the cell while making more virions, and lyse in the small intestine after uninfected *V. cholerae* have had time to replicate, providing ample hosts for subsequent rounds of phage predation. Similarly, if *V. cholerae* has bloomed in the gut followed by a subsequent expansion of the phage population to the extent that phages now outnumber hosts, it would be beneficial for ICP1 progeny phage to stay inside a cell, protected from adsorbing to cells that are already making progeny phage, and instead wait for release into the aquatic environment or ingestion by an uninfected patient via person-to-person transmission where ICP1 progeny may have better chances of finding an uninfected host.

We know that lysis timing is of paramount importance to phage fitness as evidenced by holin mutations that result in altered lysis times and lower progeny yield. The deleterious effects of accelerated lysis kinetics in the case of the S holin of λ phage, which accelerates lysis by 20 to 25 minutes and lowers progeny phage yield by two to four orders of magnitude depending on the mutation and conditions (50, 68), are obvious. Similarly, accelerated lysis by abortive infection machinery like the AbiZ protein that causes *Lactococcus lactis* infected by φ31 to lyse 15 minutes early decreases phage titers 100-fold (69). Consequently, it is no surprise that the potential benefits ICP1 sees by inhibiting lysis may not be advantageous to the phage satellite, PLE, which escapes the infected *V. cholerae* cell while protecting the *V. cholerae* population. To operate as an abortive infection mechanism, PLE must inhibit ICP1 progeny production however, we also hypothesize that PLE hijacks ICP1 structural proteins to enable its own transduction to naïve cells (23). Here, we show that the PLE collapses lysis inhibition with LidI, which is conserved as part of its anti-phage program – this adds the additional layer of complexity that is parasitizing a parasite the way a satellite parasitizes a phage. Optimal virulence is the idea that parasites must strike a balance between fitness cost to their host (virulence) and their own ability to transmit offspring to new hosts (transmission). At the limits of this theoretical trade-off are situations wherein parasites do little damage to their host over a long period of time in an effort to increase the likelihood of spreading offspring while, at the other extreme, the parasite does so much damage to the host that death may occur before the parasite can spread. In the former case, resources are left available by the parasite and these can be taken advantage of by host defense or more aggressive parasites potentially limiting the spread of the original parasite. ICP1, by nature of being a lytic phage, is a virulent parasite of *V. cholerae*, however PLE is more difficult to categorize because it is viewed as both an anti-phage mechanism that protects *V. cholerae* populations through abortive infection, while also being a parasite of ICP1 as it depends on ICP1 for horizontal spread (23). While the benefits of LIN to ICP1 have been explored in the parallel T-even phage/*E. coli* systems (10) and discussed above, PLE impact on LIN could not have been predicted – one could argue that LIN might provide PLE with more resources for horizontal transmission or counter that LIN might provide time enough for ICP1 to evade PLE-encoded anti-phage mechanisms. Indeed, initial observations of ICP1 infections of *V. cholerae* harboring PLE show accelerated lysis instead of periods of prolonged latency. Our findings suggest that the bottlenecking of the phage population (experimentally evidenced in this work by decreased phage yield in the presence of LidI and more dramatically evident in the complete block of ICP1 progeny phage production by the intact PLE) limits evolution, presumably slowing ICP1’s ability to overcome PLE. This is particularly interesting considering functionally similar <u>*S*</u>*taphylococcus* <u>*a*</u>*ureus* <u>p</u>athogenicity islands (SaPIs). SaPIs lay dormant in the chromosome much like PLE, until upon phage infection, SaPIs are induced to parasitize phage components (70). One of the many ways that PLEs and SaPIs differ is that SaPIs allow for some escape of their helper phage where as PLE completely ablates ICP1 production of progeny phage. It is easy to think of SaPIs as hitchhikers in the chromosome of the bacterial host – they integrate, take advantage of vertical transmission until the cells are challenged by the helper phage at which point they excise, inhibit phage for their own ends hijacking packaging materials, and escape the cell while allowing some progeny phage to escape with them all the while promoting diversity and horizontal gene transfer (59, 70). This ensures horizontal transfer of the SaPI as well as continued activation of SaPIs down the line by available helper phages. PLE, because it completely blocks ICP1 production acting as an abortive infection mechanism which protects *V. cholerae* at the population level, is more readily thought of as ‘on *V. cholerae’s* side’ in the dynamic evolutionary arms race. Indeed, our evidence that LidI functions through collapsing LIN supports this angle as LIN could potentially aid PLE in transduction the same way it aids ICP1 – allowing for more time to produce particles enabling transmission of PLE to larger numbers of naïve cells. There is also limited evidence that PLE uses the hijacked ICP1 machinery to transduce in nature – in the laboratory, conditions allow four of the five PLEs to integrate in many sites across the superintegron however natural isolates only ever have one of those four PLEs integrated in one specific site (23). This is indicative of vertical transmission in nature and infrequent horizontal transduction in respect to the strains we sample in epidemics, which makes it easier to reconcile PLE’s abortive infection activity.

With these evolutionary hypotheses in mind, ICP1 acquiring the CRISPR-Cas system has changed the game – single spacers encoded in ICP1 and targeting PLE can allow ICP1 to replicate while simultaneously allowing for transduction of PLE (24). If things stopped here and all spacers were singular and created equal, selection could drive PLE to act more like a typical satellite phage, embracing horizontal transfer and allowing ICP1 to slide by producing at least some progeny phage. We know, however, that this is not the case; CRISPR systems function by dynamically acquiring spacers (71) and multiple spacers can abolish PLE’s ability to transduce while also killing the cell harboring PLE, destroying any chance at horizontal or vertical transmission (24). Including this level of complication, it is no surprise PLE would employ functions like LidI to collapse lysis inhibition – why keep the cell together for long enough for rare escape phage to pick up spacers against PLE and potentially tip the balance further in the phage’s favor? It is important to continue research in this area to answer these questions that result from microbes, viruses, and satellites preying upon each other and tie these ideas together. Focusing future work on LIN and MGEs is particularly promising given that this work represents the first characterization of LIN outside of the T-even coliphages yet we found homologs of LIN machinery elsewhere. This suggests that LIN exists outside of characterized systems and, consequently, the impacts LIN are unknown and largely unexplored. In contrast, the importance of MGEs is widely accepted making the questions about the interplay between them and the complicating factors outlined here particularly attractive, especially in the context phage and phage satellites as there is increased interest in phage therapy to treat and prevent disease.

All in all, these studies characterize a novel incarnation of the classical phenomenon of LIN during phage infection of a globally relevant pathogen and a clinically relevant phage. Highlighting the phenomenon’s relevance, we found that a prevalent phage satellite encodes a conserved protein that collapses LIN, thus limiting phage yield from infection and bottlenecking the population. This reveals some of the mechanism behind the PLE’s successful anti-phage activity and lays the groundwork for asking classic questions about evolutionary theory in respect to optimal virulence and selective pressures in complicated parasitic relationships, all the while providing insight into an ongoing evolutionary arms race that impacts human health.

## Materials and Methods

### Resource Table

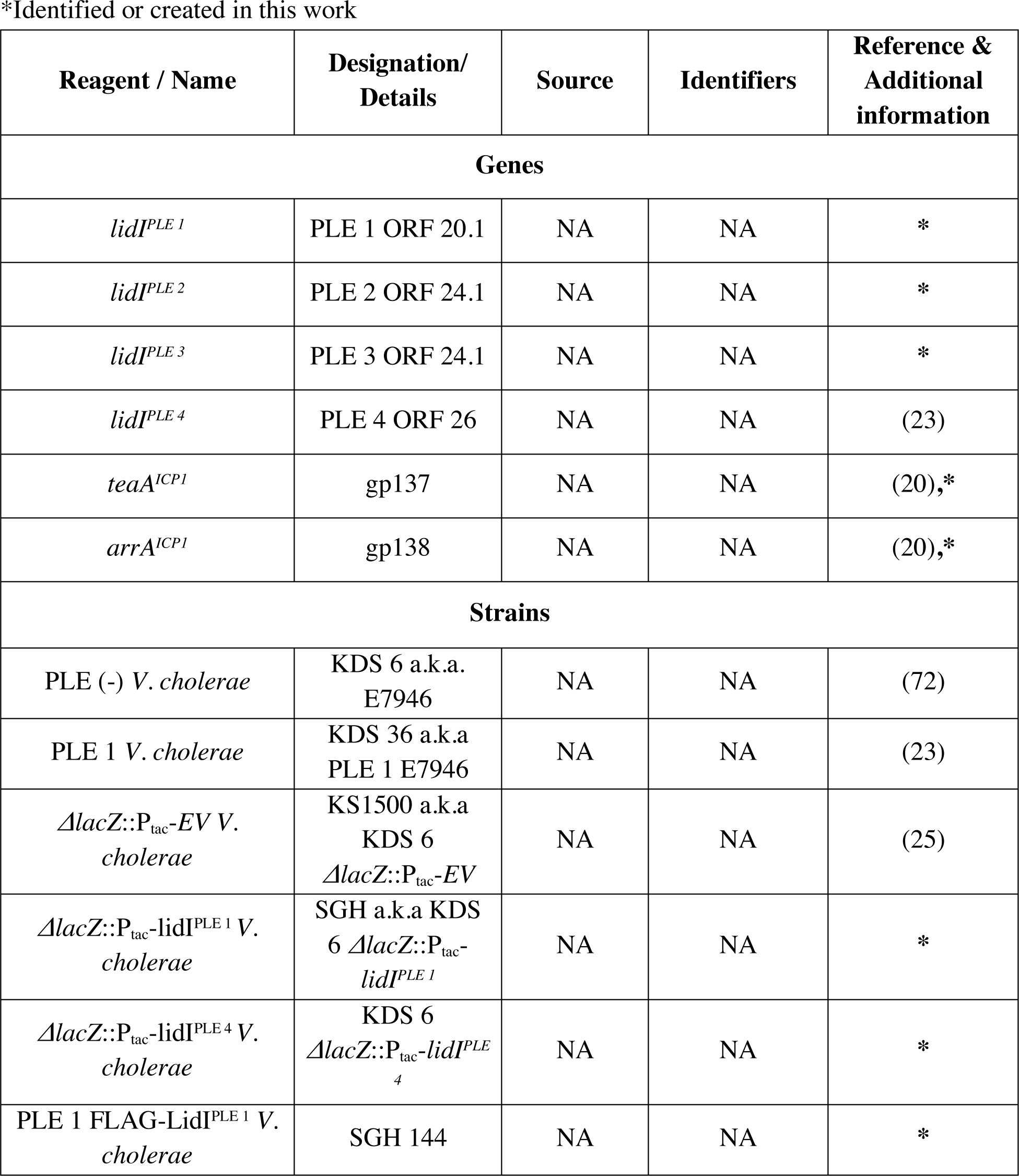
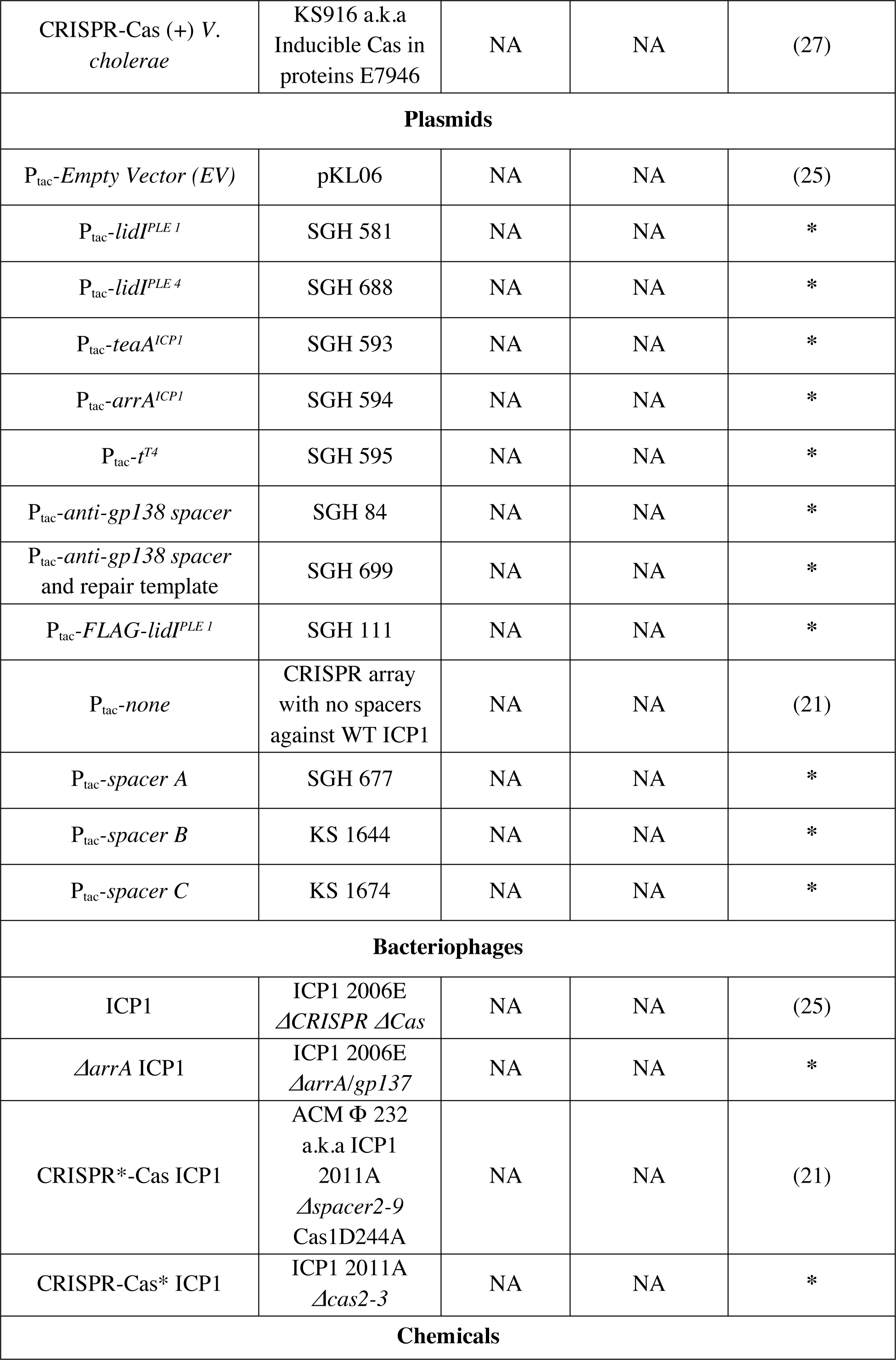
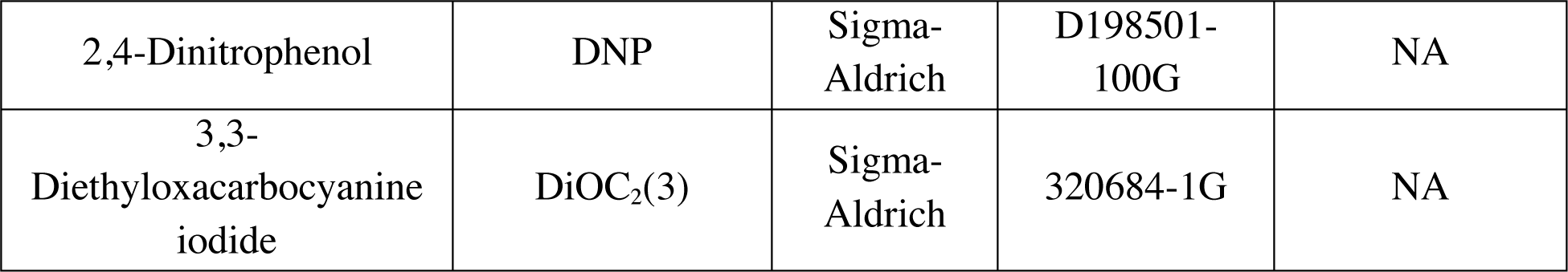

| Reagent / Name | Designation/<br>Details | Source | Identifiers | Reference &<br>Additional<br>information |
| --- | --- | --- | --- | --- |
| **Genes** |  |  |  |  |
| *lidI<sup>PLE 1</sup>* | PLE 1 ORF 20.1 | NA | NA | * |
| *lidI<sup>PLE 2</sup>* | PLE 2 ORF 24.1 | NA | NA | * |
| *lidI<sup>PLE 3</sup>* | PLE 3 ORF 24.1 | NA | NA | * |
| *lidI<sup>PLE 4</sup>* | PLE 4 ORF 26 | NA | NA | (23) |
| *teaA<sup>ICP1</sup>* | gp137 | NA | NA | (20),* |
| *arrA<sup>ICP1</sup>* | gp138 | NA | NA | (20),* |
| **Strains** |  |  |  |  |
| PLE (-) *V. cholerae* | KDS 6 a.k.a.<br>E7946 | NA | NA | (72) |
| PLE 1 *V. cholerae* | KDS 36 a.k.a<br>PLE 1 E7946 | NA | NA | (23) |
| $\Delta lacZ::P_{tac}-EV$ *V. cholerae* | KS1500 a.k.a<br>KDS 6<br>$\Delta lacZ::P_{tac}-EV$ | NA | NA | (25) |
| $\Delta lacZ::P_{tac}-lidI^{PLE 1}$ *V. cholerae* | SGH a.k.a KDS<br>6 $\Delta lacZ::P_{tac}-$<br>*lidI<sup>PLE 1</sup>* | NA | NA | * |
| $\Delta lacZ::P_{tac}-lidI^{PLE 4}$ *V. cholerae* | KDS 6<br>$\Delta lacZ::P_{tac}-lidI^{PLE 4}$ | NA | NA | * |
| PLE 1 FLAG-LidI<sup>PLE 1</sup> *V. cholerae* | SGH 144 | NA | NA | * |
| CRISPR-Cas (+) *V. cholerae* | KS916 a.k.a Inducible Cas in proteins E7946 | NA | NA | (27) |
| **Plasmids** |  |  |  |  |
| $P_{tac}$-Empty Vector (EV) | pKL06 | NA | NA | (25) |
| $P_{tac}$-*lidI*<sup>PLE 1</sup> | SGH 581 | NA | NA | * |
| $P_{tac}$-*lidI*<sup>PLE 4</sup> | SGH 688 | NA | NA | * |
| $P_{tac}$-*teaA*<sup>ICP1</sup> | SGH 593 | NA | NA | * |
| $P_{tac}$-*arrA*<sup>ICP1</sup> | SGH 594 | NA | NA | * |
| $P_{tac}$-*t*<sup>T4</sup> | SGH 595 | NA | NA | * |
| $P_{tac}$-*anti-gp138 spacer* | SGH 84 | NA | NA | * |
| $P_{tac}$-*anti-gp138 spacer* and repair template | SGH 699 | NA | NA | * |
| $P_{tac}$-FLAG-*lidI*<sup>PLE 1</sup> | SGH 111 | NA | NA | * |
| $P_{tac}$-none | CRISPR array with no spacers against WT ICP1 | NA | NA | (21) |
| $P_{tac}$-*spacer A* | SGH 677 | NA | NA | * |
| $P_{tac}$-*spacer B* | KS 1644 | NA | NA | * |
| $P_{tac}$-*spacer C* | KS 1674 | NA | NA | * |
| **Bacteriophages** |  |  |  |  |
| ICP1 | ICP1 2006E ΔCRISPR ΔCas | NA | NA | (25) |
| ΔarrA ICP1 | ICP1 2006E ΔarrA/*gp137* | NA | NA | * |
| CRISPR*-Cas ICP1 | ACM $\Phi$ 232 a.k.a ICP1 2011A Δspacer2-9 Cas1D244A | NA | NA | (21) |
| CRISPR-Cas* ICP1 | ICP1 2011A Δcas2-3 | NA | NA | * |
| **Chemicals** |  |  |  |  |
| 2,4-Dinitrophenol | DNP | Sigma-Aldrich | D198501-100G | NA |
| 3,3-Diethyloxacarbocyanine iodide | DiOC<sub>2</sub>(3) | Sigma-Aldrich | 320684-1G | NA |

### Bacteria and phage propagation

Bacteria were propagated at 37°C via streaking from frozen glycerol stocks on solid LB agar plates and growth in Miller LB (Fisher Bioreagents) with aeration. Media was supplemented with chloramphenicol (2.5 μg/mL for *V. cholerae* and 25 μg/mL chloramphenicol for *E. coli*), kanamycin (75 μg/mL), ampicillin (100 μg/mL), and streptomycin (100 μg/mL) when appropriate. Cell densities were measured at OD_600_ in tubes (Biochrom Ultrospec 10; 10 mm pathlength referred to as OD_600-tube_) and in 96-well plates (all reported OD_600_ measurements in figures have a pathlength equivalent to 150 μL in Costar Clear 96-well plates (Corning)). To induce chromosomal constructs in both liquid and top agar, plasmid constructs in top agar, and the plasmid P_tac_-*arrA* construct for complementation, 1 μM isopropyl β-D-1-thiogalactopyranoside (IPTG) and 1.5 μM theophylline were added to cultures while the remaining plasmid constructs were induced with 0.125 μM IPTG and 0.1875 μM theophylline in liquid cultures. Plasmid constructs were used for all experiments other than phage infection yield experiments which utilized chromosomal constructs to decrease leaky expression in uninduced strains.

Bacteriophages were propagated on PLE (-) *V. cholerae* hosts and prepped via polyethylene glycerol precipitation or concentration and media exchange on Amicon Ultra – 15 (Millipore) centrifugal filters (73, 74). Stocks were stored in sodium chloride-tris-ethylenediaminetetraacetic acid buffer (STE), and quantified via the soft agar overlay method (73). Briefly, titering was completed by growing *V. cholerae* to mid-log, infecting with cultures with diluted phage, and allowing adsorption to occur for 7 to 10 minutes before plating on 0.5% LB top agar. Mechanical lysis of cultures infected with phage was accomplished by mixing chloroform into cultures then letting cultures stand at room temperature for ten minutes before spinning at 5,000 × G for 15 minutes at 4°C and removing the supernatant for further analysis.

### Cloning and strain production

Chromosomal integrations in *V. cholerae* were accomplished through natural transformation of linear DNA created via splicing by overlap extension PCR (75). In the case of deletions, antibiotic resistance cassettes were integrated into the locus of the deleted gene and subsequently flipped out as previously described (76). Plasmids were constructed with Gibson Assembly and Golden Gate reactions. Phage mutants were selected as previously described (27). Briefly, complementary spacer oligos were annealed and inserted into plasmid-borne CRISPR array. This plasmid was mated into a strain of *V. cholerae* engineered to include an inducible Type 1-E CRISPR-Cas system (CRISPR-Cas (+) *V.* cholerae). This system in the host strain was induced for 20 minutes before ICP1 infection for plaque assays on 0.5% LB top agar containing antibiotics to maintain the CRISPR array plasmid. Plaques were picked into STE, purified on the same host twice, and genomic DNA was prepped for PCRs with the DNeasy Blood and Tissue Kit (Qiagen). Phages were subjected to PCR of the targeted gene and subsequent Sanger sequencing. Clean knockouts were accomplished by adding a repair template of homologous sequence containing the desired deletion and flanking DNA to the plasmid containing the CRISPR array as previously described (27).

### Lysis kinetics

*V. cholerae* strains were grown in 2 mL cultures to an OD_600(tube)_=0.3 and 150 μL of cultures were added to 96-well plates. Inducers and phage were pre-aliquoted in plates unless otherwise specified. OD_600_ within the plate was read for each sample on the SpectraMax i3x (Molecular Devices) plate reader every two minutes with one minute of shaking between each read while the machine incubated cultures at 37°C. Assays were interrupted for the addition of phage, inducers, DNP (2 μL of 8.4 mM DNP dissolved in 80% ethanol for a final concentration of 110 μM DNP), or ethanol (2 μL of 80% ethanol) during which samples were removed from the plate reader briefly before measurement was resumed. This enabled superinfection of cultures (initial cultures were infected with ICP1 MOI=1, returned to the plate reader for four minutes, and subsequently superinfected with ICP1 MOSI=5), addition of DNP in ethanol or ethanol alone to ICP1 MOI=5 infected cultures 25 minutes post-infection, and DNP addition to induced cultures after 20 minutes of growth with the inducers.

### Phage yield from high MOI infection determination

For each strain, three 2 mL cultures of *V. cholerae* were grown. The first was grown to an OD_600-tube_=0.15, inducer was added, and the culture was returned to the incubator; this culture served as the pre-induced culture. All cultures were grown for an additional 20 minutes to OD_600-tube_=0.3 at which point ICP1 MOI=5 was added. Inducer was added to one tube at this time; this culture served as the culture induced at time zero. Tubes were returned to the incubator for 5 minutes for phage adsorption. Cultures were spun at 5,000 × G for 3 minutes to pellet cells. Unadsorbed phage was aspirated off and cells were washed once with 1 mL of prewarmed LB with or without inducer. Cells were spun again and resuspended in media with and without inducer at which point OD_600-tube_ was determined. Cultures (150 μL) were moved to 96-well plates with one well designated to the pre-induced culture, one well devoted to the culture induced at time zero, and three wells filled with uninduced culture. The plate was returned to the 37°C incubator to shake at 230 RPM. Inducer was added to two wells of the uninduced cultures 20 and 40 minutes post-infection respectively, leaving one uninduced control. The experiment was ended with mechanical lysis of cultures and subsequent quantification of phage titers.

### PMF determination

*V. cholerae* containing the specified plasmids were grown to OD_600-tube_=0.2. Inducers (125 μM IPTG and 187.5 μM) were added and cells were grown at 37°C with aeration for 30 minutes before 0.5 mL were pelleted at 5,000 × G for 3 minutes and resuspended in 0.1 mL of phosphate buffered saline (pH 7.2; Gibco Life Technologies) containing 20 μM 3,3’-diethloxacarbocyanine iodide (DiOC_2_(3); Sigma-Aldrich). Fluorescence measurements were completed in black 96-well half-volume plates (Corning) in the SpectraMax i3x (Molecular Devices) with 480(508) and 488(650) excitation(emission) wavelength settings.

### Plaque analysis

*V. cholerae* strains were grown to mid-log before plaquing assays. For EOP experiments, plasmid constructs were induced for 20 minutes prior to infecting and plating with antibiotic and inducer in the subsequent 0.5% LB top agar. For plaque edge and area analysis, plasmids were not induced prior to plating on 0.5% and 0.3% top agar, respectively, containing antibiotics and inducer. Plates solidified at room temperature prior to incubation at 37°C. For spot plates, *V. cholerae* was mixed with 0.5% top agar prior to infection, vortexed, and poured onto an LB agar plate to solidify before 3 μL spots of phage dilutions were overlaid on the agar. Plates were allowed to dry prior to incubation at 37°C. To visualize plaques, plates were scanned on the EPSON Perfection V800 Dual Lens scanner. Area of plaques were calculated via Fiji (77) with particle analysis of isolated singular plaques.

### Western blots

Plasmid EV and FLAG-LidI^PLE 1^ were grown to OD_600-tube_=0.2 then induced with 1 mM IPTG and 1.5 mM for 40 minutes before 0.5 mL samples were taken. To observe expression during infection, strains were grown to OD_600-tube_=0.3, infected with ICP1 MOI=2 with 4 mL samples taken at the labeled timepoints. Samples were prepared and visualized as previously described (25). Briefly, samples were mixed with one volume of cold methanol, pelleted at 15,000 × G at 4°C for 15 minutes, washed with 1 mL cold PBS, and pelleted. Pellets were resuspended in PBS with XT sample buffer and reducing agent (Bio-Rad), vortexed, and boiled for 10 minutes before being run on 4-12% Bis-Tris SDS gels (Bio-Rad Criterion XT). Gels were transferred via the Trans-Blot Turbo (Bio-Rad) and visualized with rabbit-α-FLAG (1:3,000) primary and goat-α-rabbit-HRP conjugated secondary antibodies on the ChemiDoc MP Imaging System (Bio-Rad).

### CRISPR-Cas targeting of phage populations

*V. cholerae* harboring EV or LidI^PLE 1^ plasmids were grown to OD_600-tube_=0.15 before being induced and returned to the incubator to grow with aeration until OD_600-tube_=0.3. At this point 150 μL of culture were added to 96-well plates and infected with ICP1 MOI=0.1. After lysis 90 minutes post-infection, any remaining cells were mechanically lysed and the resulting phage population was plaqued on CRISPR-Cas (+) *V. cholerae* as previously described (27). Briefly, Cas (+) *V. cholerae* harboring CRISPR array plasmids were induced 20 minutes before phage populations were titered on each specified *V. cholerae* strain. EOPs were determined by dividing the number of PFU/mL on *V. cholerae* containing spacers by the PFU/mL on the CRISPR-Cas (+) *V. cholerae* that did not contain a spacer against the phage (denoted as spacer ‘none’).

### Coinfection for homologous recombination

Coinfection experiments were completed in the same manner as the CRISPR-Cas targeting of phage populations described above with minor alterations: instead of infection with WT ICP1, cultures were coinfected in the plate with CRISPR*-Cas ICP1 and CRISPR-Cas* ICP1 each at an MOI=0.01, observed for 200 minutes in the plate reader, and, after mechanical lysis, phage populations were plaqued on PLE (-) *V. cholerae* and PLE 1 *V. cholerae.* Control infections with only one phage never formed plaques on PLE 1 *V. cholerae.* The proportion of phages that successfully recombined was determined by the dividing the PFU/mL of each phage population on PLE 1 *V. cholerae* divided by the PFU/mL on PLE (-) *V. cholerae*.

### Bioinformatics

Transmembrane domains were predicted by conversion of all predicted open reading frames to amino acid sequence by CLC (78) before analysis with TMHMM Server v. 2.0 (49). PRALINE was used to create amino acid alignments of LidI (79). Homologs of TeaA (30% identity over 85% of the query) and ArrA (20% identity over 75% of the query) were identified with BLASTP (80) and arranged into phylogenetic trees as previously described (22). Briefly, alignments were completed with MUSCLE v3.8.31 (81) and a bootstrapped (n=100) maximum-likelihood phylogenic tree was solved with PhyML 3.0 (82). Default settings were used for amino acid sequences: automatic model selection with Akaike Information Criterion; SPR tree improvement with n=10 random starting trees. Trees were visualized with FigTree (83).

## Supplemental Material

Supplemental figures are included following the references. Supplemental material for this article may be found by emailing the corresponding author after the paper has been peer reviewed.

## Acknowledgements

This work was supported by the National Institute of Allergy and Infectious Diseases [R01AI127652 to K.D.S.]; K.D.S. is a Chan Zuckerberg Biohub Investigator and holds an Investigators in the Pathogenesis of Infectious Disease Award from the Burroughs Wellcome Fund. Thanks to the Seed Lab for thoughtful discussions and help with special thanks to Zoe Netter, Caroline Boyd, Zach Barth, and Amelia McKitterick for reviewing drafts.

## Supplemental Material

**Supplemental Fig. 1.**
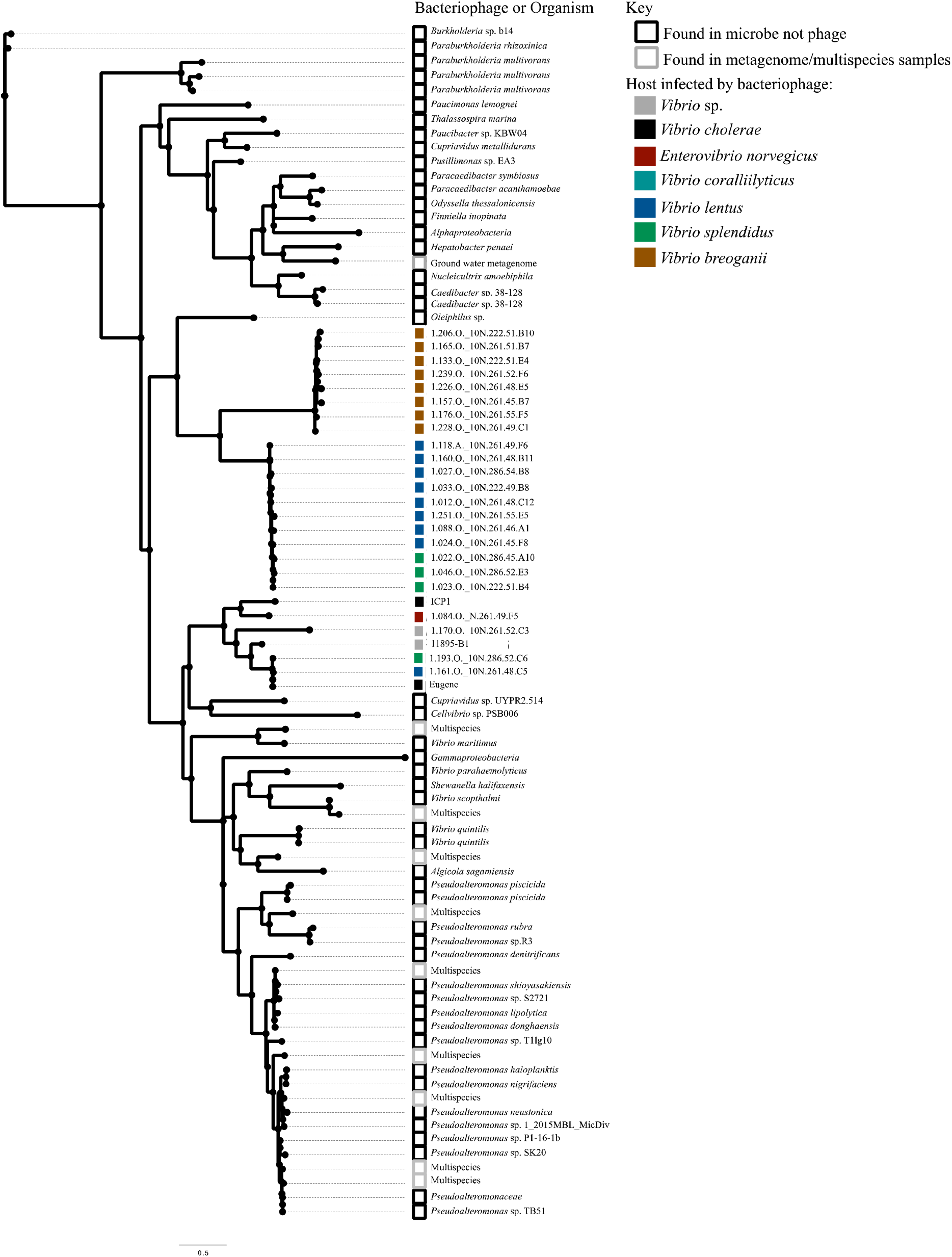
Proteins with similarity to TeaA. This unrooted tree shows proteins identified with BLASTP that share 30% identity with TeaA over 85% of the query. The source of the sequence is shown in the label on the tree while colored blocks denote the host the phage infects or whether the sequence was found in a microbe or multispecies sequence entry.

**Supplemental Fig. 2.**
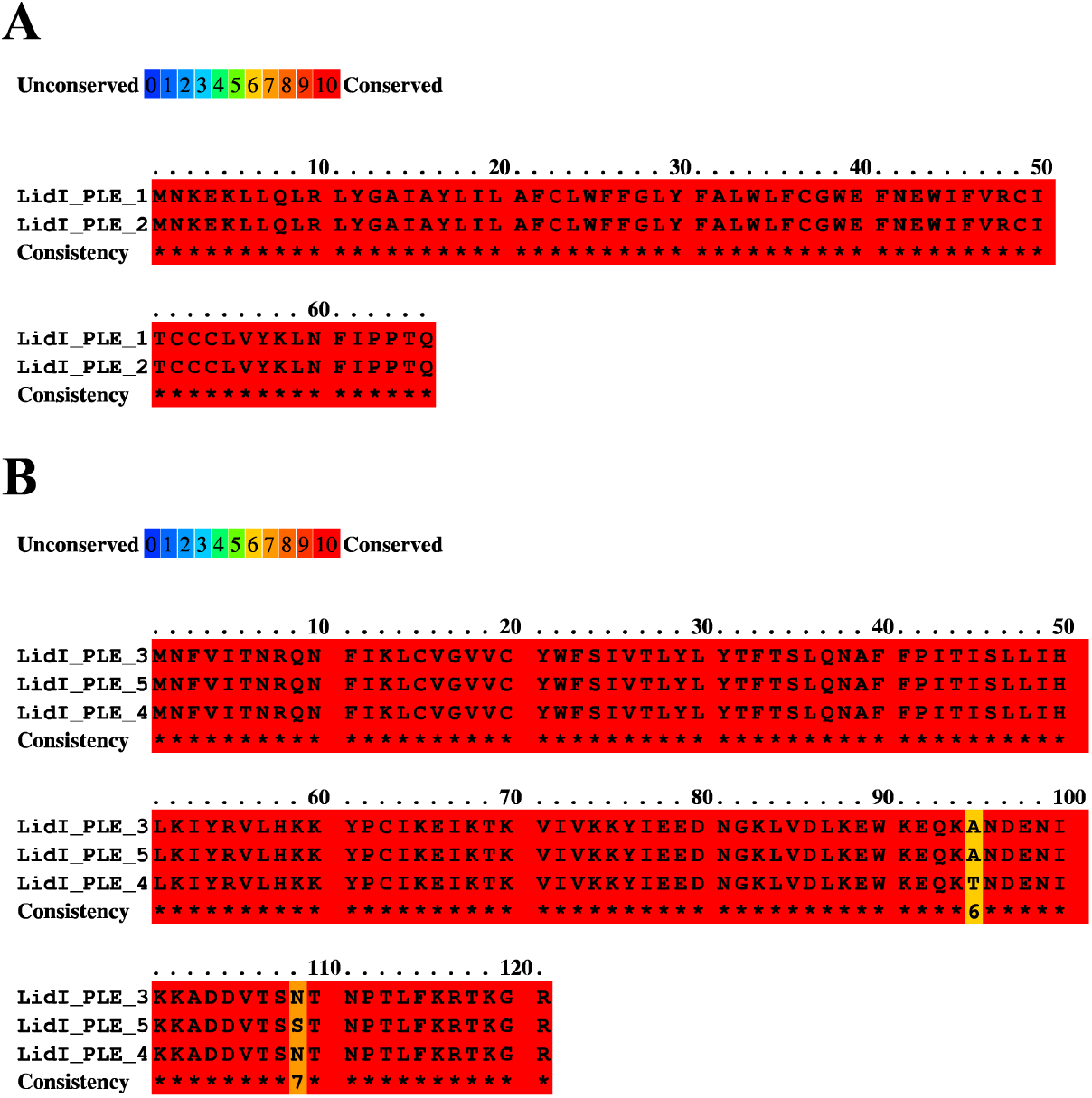
PRALINE alignment of LidI homologs. LidI homologs cluster neatly in two groups: **A** LidI^PLE 1^ and LidI^PLE 2^ and **B** LidI^PLE 3^, LidI^PLE 4^, and LidI^PLE 5^

**Supplemental Figure 3.**
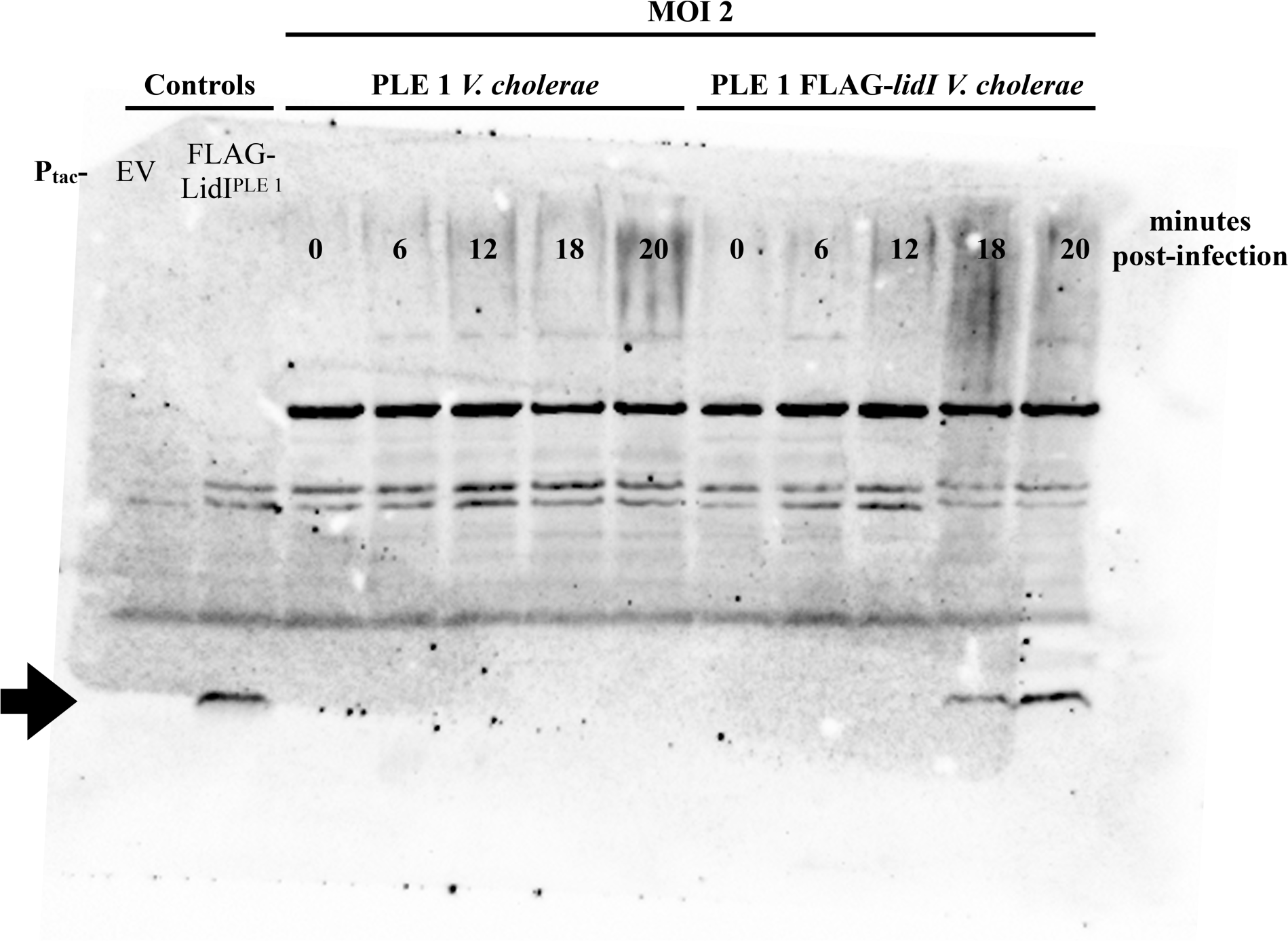
Complete Western Blot. Tagged LidI^PLE 1^ was visualized via western blot. EV and FLAG-LidI^PLE 1^ expressed *in trans* served as controls showing FLAG-LidI^PLE 1^ specificity as well as the presence of background bands. In PLE 1 *V. cholerae*, there is no detectable band similar in size to tagged LidI^PLE 1^ (as indicated by the arrow). When LidI^PLE 1^ is tagged in the native locus and strains are infected with ICP1, FLAG-LidI^PLE 1^ is visible late in infection (18 and 20 minutes post-infection).

**Supplemental Figure 4.**
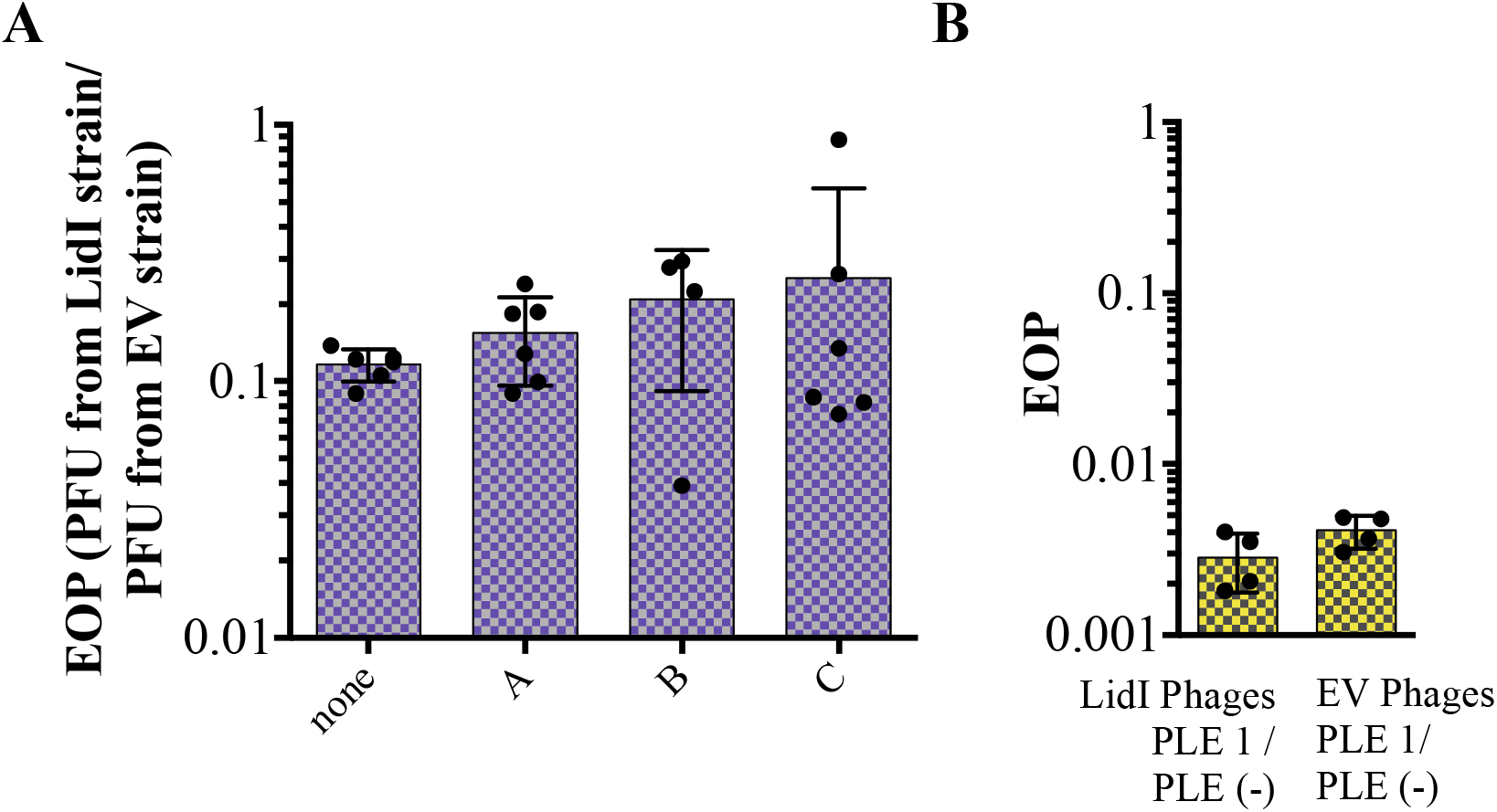
EOP of evolution experiments in the presence of LidI^PLE 1^. **A** EOP was determined between PFUs from LidI^PLE 1^ *V. cholerae* infections and PFUs from EV *V. cholerae* infections plaqued on CRISPR-Cas (+) *V. cholerae* harboring each of the specified spacers. **B** EOP was determined between PFUs of a population plaqued on PLE 1 *V. cholerae* in comparison to the PFUs of the same population on PLE (-) *V. cholerae*.

